# Optimizing light traps for littoral mysids and mesopredatory fish in the Baltic Sea: environmental drivers and seasonal monitoring efficacy

**DOI:** 10.64898/2026.07.01.735747

**Authors:** Martin Ogonowski

## Abstract

Littoral mysids facilitate benthic-pelagic coupling through horizontal migration, yet quantitative monitoring in structurally complex habitats remains methodologically challenged where traditional active gears fail. We evaluated the efficacy of standardized light traps for monitoring littoral mysids (*Neomysis integer*, *Praunus flexuosus*) and mesopredatory three-spined sticklebacks (*Gasterosteus aculeatus*) in the northern Baltic proper, Baltic Sea. Using a paired experimental design with predator-exclusion and unmodified traps, alongside concurrent passive benthic trapping, we assessed abiotic drivers affecting catchability, biotic interactions, and statistical power to monitor changes in population size over time. Results indicated significant biotic interference: unmodified traps attracted high densities of sticklebacks, which reduced mysid catches by approximately 85% through predation or behavioural avoidance. Consequently, physical predator exclusion is mandatory for accurate mysid sampling. Generalized Linear Mixed Models (GLMMs) confirmed that catch rates for all taxa were primarily driven by night duration rather than water temperature. While passive benthic trap catches tracked metabolic activity (peaking in warm summer months), light trap efficiency peaked in spring and collapsed during summer, confirming that sampling efficiency was strictly limited by the short duration of the night. Simulation-based power analysis revealed a stark contrast in monitoring utility based on spatial aggregation. For highly aggregated mysids, the method demonstrated low precision (Power < 0.25 to detect a 50% decline), rendering it suitable primarily for detecting substantial population collapses (>90%). In contrast, for less aggregated sticklebacks, the method achieved a more robust statistical power (>0.80 for a 60% decline), validating light traps as a precise tool for monitoring these abundant mesopredators. We conclude that light traps fill a critical methodological gap for winter and early spring monitoring when traditional passive gears underperform. Appropriate abundance indices should be based on statistical models accounting for night duration and strictly employ physical exclusion barriers when targeting mysids.

## 2 Introduction

In coastal ecosystems, littoral zones function as critical nurseries and feeding grounds, supporting high biodiversity and biomass. Within these structurally complex habitats, littoral mysids (e.g., *Neomysis integer*, *Praunus flexuosus*) play a pivotal but often unquantified ecological role. Unlike their pelagic counterparts that couple benthic and surface waters through deep diel vertical migration (Rudstam *et al*. 1989; Boscarino *et al*. 2009; Ogonowski, Hansson and Duberg 2013), littoral mysids facilitate benthic-pelagic coupling through diel (Debus, Mehner and Thiel 1992) and seasonal horizontal shifts (Kotta and Kotta 1999; Lesutienė *et al*. 2008). During the day, they utilize nearshore vegetation and rocky substrates as refugia from visual predators; at night, they migrate horizontally into open water to forage on zooplankton (Jumars 2007). This behaviour enables the transfer of carbon from nearshore production to higher trophic levels, including commercially and ecologically important fish species such as European perch (*Perca fluviatilis*) and three-spine sticklebacks (*Gasterosteus aculeatus*).

Despite their abundance and functional importance, littoral mysids are frequently underrepresented in environmental monitoring programs and food-web models. Their exclusion is largely attributable to the difficulty of reliably quantifying their abundance in free living populations (Jumars 2007). Traditional active sampling methods, such as drop frames and seine nets, are often rendered inoperable by the physical complexity of rocky substrates or dense vegetation. Conversely, hydroacoustic methods, effective for pelagic aggregations, cannot resolve targets against the high acoustic backscatter of the benthos. Consequently, determining the true contribution of mysids to ecosystem functioning requires standardized sampling methods that can function effectively within complex habitats while retaining a high catchability.

An alternative passive method is the use of light traps which commonly are used for biodiversity monitoring of larval fishes (Doherty 1987; Pierce *et al*. 2006; Reynalte-Tataje *et al*. 2024), mapping distribution patterns of pelagic lobster larvae (Øresland 2008; Sigurdsson, Morse and Rochette 2014) and early-warning detection of invasive invertebrates such as the bloody red mysid *Hemimysis anomala* (Brown *et al*. 2017), though their application for quantitative monitoring remains underutilized. Consequently, the performance of light traps in complex, vegetated habitats, where biotic interactions such as predation may severely compromise catch efficiency, has not been rigorously evaluated for native mysid species like *Neomysis integer*. Mysids exhibit strong phototactic responses, varying by species and illumination intensity (Gal *et al*. 1999; Viherluoto and Viitasalo 2001). Although light traps act as non-specific samplers and may, like most other trap designs be biased by “in-trap” predation, they offer distinct advantages: they are effective for phototactic species, cause low environmental disturbance, and allow for sampling in sensitive, structurally complex areas over extended periods (Brogan 1994; Meekan *et al*. 2001; Porter 2016). Despite this potential, a standardized protocol for their use in marine and brackish monitoring remains undeveloped (McLeod and Costello 2017).

The Baltic Sea offers an ideal system to evaluate these sampling methods due to its low species diversity and low functional redundancy (Elmgren and Hill 1997). This simplicity aids in interpreting food web interactions, particularly in the face of anthropogenic pressures. Baltic coastal ecosystems are currently undergoing large-scale ecological changes, notably the decline of large predatory fish such as northern pike and a concurrent increase in mesopredatory fish populations, specifically the three-spined stickleback (Bergström *et al*. 2015, 2022; Olsson *et al*. 2023). This population expansion has been implicated in a trophic cascade where increased predation on grazers releases filamentous algae from control, degrading essential nursery habitats (Donadi *et al*. 2017; Eklöf *et al*. 2020).

The effective recovery of predatory fish requires disentangling these complex food web interactions. Sticklebacks are known to prey on pike larvae (Nilsson, Flink and Tibblin 2019), but the role of other organisms in this predator-prey reversal remains unquantified. Mysids, being effective zooplankton predators, potentially compete for resources with both sticklebacks and first-feeding pike larvae. However, without a reliable method to monitor mysids in the complex littoral vegetation where these interactions occur, their contribution to this ecosystem-change cannot be assessed.

The aim of this study was to describe and evaluate the efficacy of light traps for monitoring littoral mysids in complex coastal environments, focusing on two common Baltic species, *Neomysis integer* and *Praunus flexuosus*. We assessed potential biotic and abiotic factors affecting catch rates to develop a standardized catch-per-unit-effort (CPUE) index suitable for monitoring population trends. Additionally, we investigated the effectiveness of light traps in capturing fish mesopredators (sticklebacks) to determine if this method can simultaneously monitor interacting guild members in these changing coastal ecosystems.

## 3 Methods

### 3.1 Study area

Field trials were conducted in four coastal bays at Askö, an island in the southern Stockholm archipelago, Sweden (58° 49.015’ N, 17° 39.243’ E, Figure 1). These bays are representative of the region’s littoral habitats, which are increasingly dominated by mesopredatory sticklebacks following regional declines in larger predatory fish (Eklöf *et al*. 2020; Olsson *et al*. 2023). The bays feature mixed substrates (sand, mud, and gravel) with varying coverage of macroalgae (e.g., *Fucus vesiculosus*) and vascular plants. Field operations were facilitated by the infrastructure and boating support provided by the Askö Laboratory (Stockholm University Marine Research Centre).

**Figure 1.**
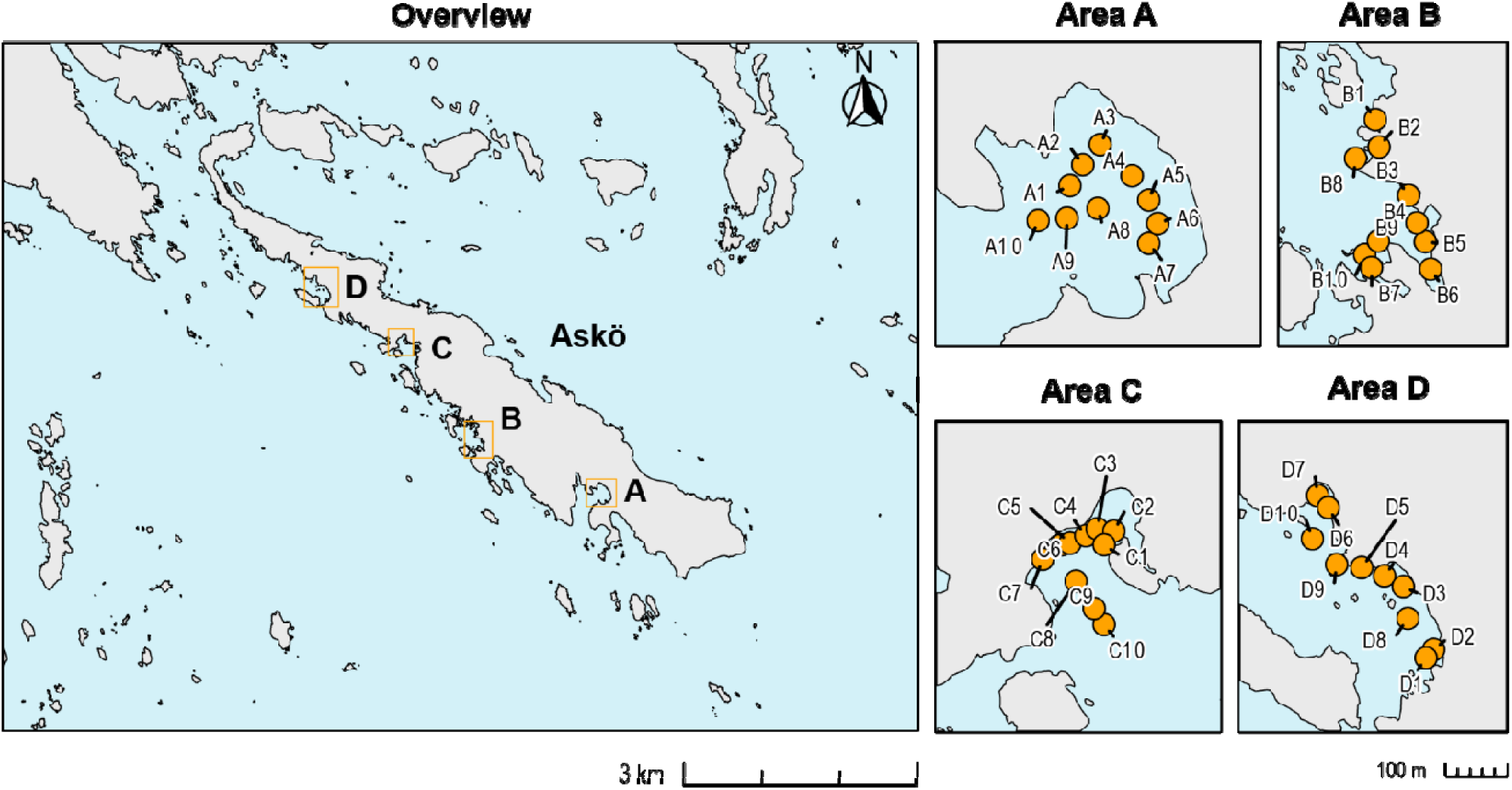
Location of sampling stations (Areas A-C) in four shallow bays on Askö island, Southern Stockholm archipelago, Sweden. Ten mooring stations for light traps and benthic traps were distributed within each area amounting to a total of 40 moorings.

### 3.2 Experimental treatments

We utilized a quatrefoil light trap design with green LED-illumination (after Floyd, Courtenay and Hoyt 1984) targeting mysids and sticklebacks (Figure 2). A preliminary pilot study indicated that the standard trap design attracted high numbers of sticklebacks (*G. aculeatus* and *Pungitius pungitius*), which appeared to reduce mysid capture rates through avoidance behaviour (Lindén, Lehtiniemi and Viitasalo 2003; Boscarino *et al*. 2007; Barrios-O’Neill *et al*. 2014) or “in-trap” predation (Vilizzi *et al*. 2008). To assess this effect, we deployed the traps in two configurations.

**Figure 2.**
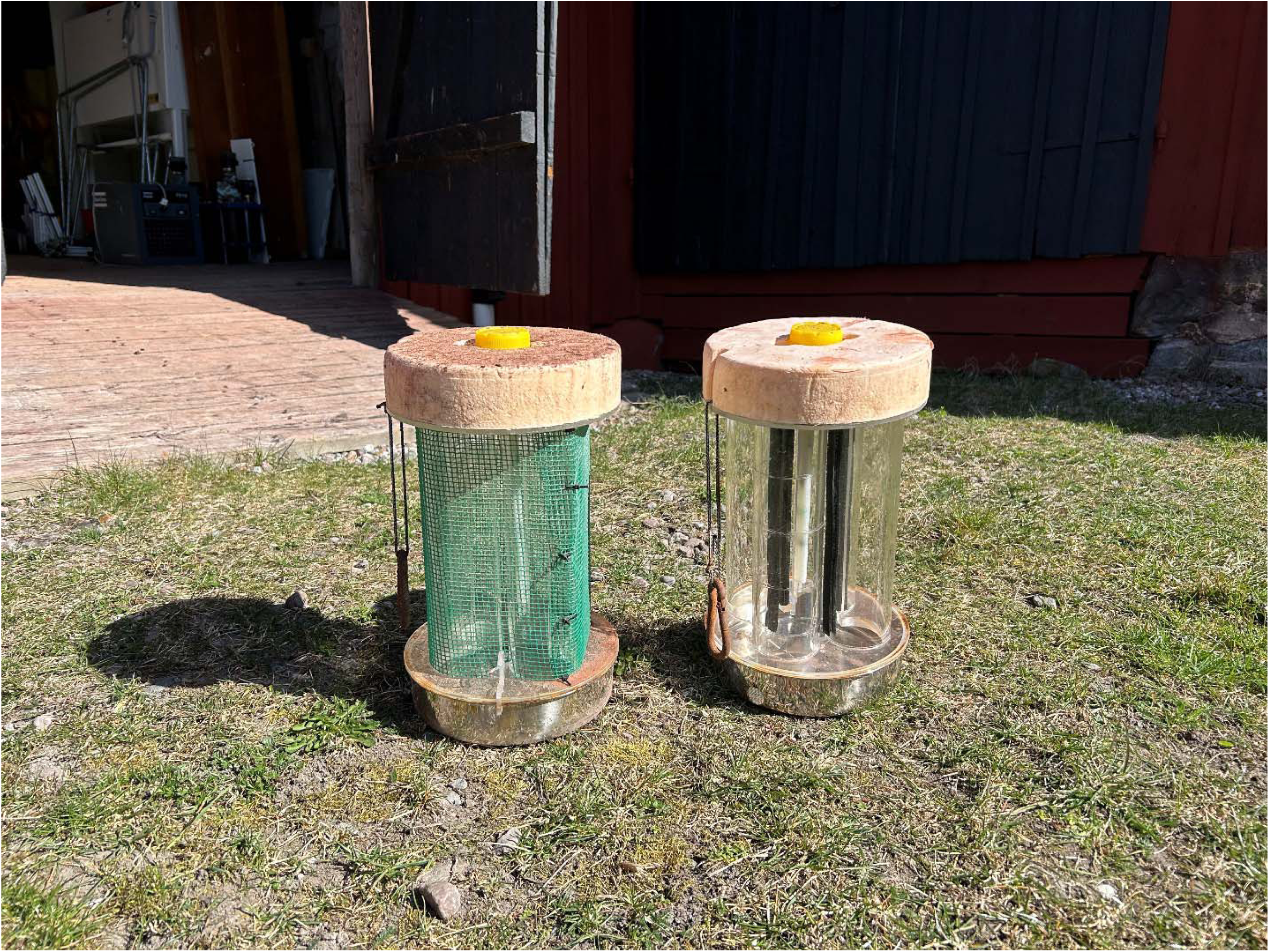
Modified (left) and Unmodified (right) quatrefoil light traps. The modified trap has a protective green netting to keep potential predators away. The unmodified trap has a flexible U-channel edge trim (black strips) applied to one side of the entrance slits to decrease the gape width of the trap.

The first, basic configuration represented the standard quatrefoil design and is hereafter referred to as the “Unmodified trap design”. Due to minor imprecision in the machining process, the initial slit widths exceeded the target specifications. To correct this, a flexible U-channel edge trim (PVC with internal steel reinforcement; 9 mm width, Biltema, Sweden) was mounted along one vertical edge of each entrance slit. This reduced the effective entrance width to the required 5–7 mm.

The second configuration included a modification to physically exclude sticklebacks while allowing mysid entry (cf. Vilizzi *et al*. 2008; Powell and Mandrak 2025). This modification (hereafter called the “Modified trap design”) consisted of a green plastic garden net (2 × 2 mm mesh) which was mounted over the entrance slits using cable ties. The U-channel edge trim was not applied to these traps, as the mesh defined the effective entrance size (Figure 2).

Because light traps are size-selective and rely on phototaxis, we used collapsible benthic traps to assess the ambient population structure of sticklebacks independent of light attraction. Although originally designed for freshwater crayfish (Kayoba, Skara, Sweden), these traps effectively sample sticklebacks (Donadi *et al*. 2020). The traps measured 45 × 24 × 24cm with a 3 mm mesh size and featured two 5.5 cm diameter circular entrances. Details on the trap design and illumination configuration are provided in the Supporting Information, sections S1-S3.

### 3.3 Deployment and sampling routine

Sampling stations were established using a single mooring line anchored by a 4 kg weight. The light trap was attached to a surface buoy, floating on the surface. The benthic trap was attached to the same mooring line but positioned on the seafloor (Figure 3). This paired arrangement allowed for the assessment of littoral mysid abundance and the effect of light on stickleback catch rates.

**Figure 3.**
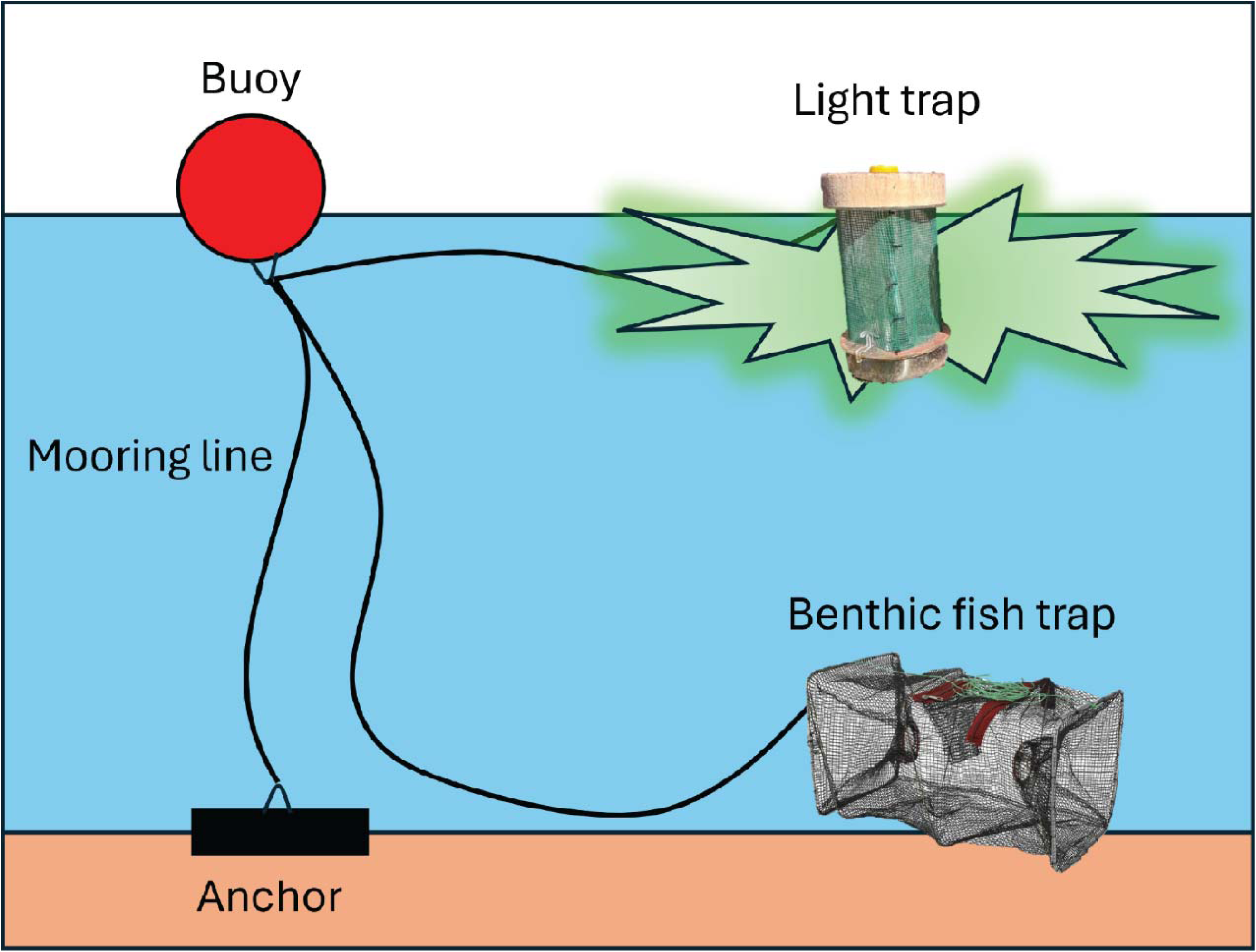
Schematic image of the trap mooring.

A systematic balanced and stratified design was employed within each of four areas (coastal bays) to cover the littoral gradient (Stevens Jr and Olsen 2003; Perret *et al*. 2022). Ten traps were deployed per area per sampling event: seven stations were distributed along the vegetated shoreline (approx. 0.5 m depth), and three stations were positioned further offshore (0.5–1.5 m depth). To ensure balanced coverage of trap types across depths and dates, Modified and Unmodified traps were deployed in an alternating sequence at each depth stratum (n = 5 Modified and n = 5 Unmodified traps per area per sampling event).

Sampling was conducted weekly between 25 April and 19 June 2023, with one additional follow-up survey on 10 July (n = 11 sampling events). Traps were deployed from a small boat with a mean soak time of 24 h (± 2.1 h SD).

### 3.4 Sample collection and preservation

Upon retrieval, light traps were lifted vertically, allowing the water column to drain through the mesh-covered aperture of the collection vessel. The catch (mysids and fish) was immediately transferred to plastic containers.

All sampling, handling, and euthanasia protocols for vertebrate species (predominantly *G. aculeatus*) were explicitly reviewed and approved by the Linköping Animal Experimental Ethics Committee under permit reference number Dnr 23379-2022. To ensure rapid euthanasia and preservation as stipulated and approved under this permit, a two-step method was utilized. The catch was immersed in an initial concentration of approximately 70% ethanol (achieved by adding 95% ethanol to the sample water) to induce rapid cessation of biological activity. Following this, the sample was transferred to fresh 70% ethanol for long-term preservation and to minimize tissue shrinkage or distortion.

Catches from benthic traps were treated in a similar fashion; however, because benthic hauls could be substantial, we only euthanized and preserved a randomized subsample (approximately 50 *G. aculeatus* individuals per sampling area and sampling event) for downstream size measurements, releasing the remaining catch at the site of capture.

### 3.5 Environmental variables

Concurrent with trap retrieval, physicochemical parameters were recorded using a Rinko ASTD-102 profiler (JFE Advantech Co., Ltd., Japan). Measured variables included water temperature, salinity, dissolved oxygen, chlorophyll-*a* fluorescence, turbidity, and conductivity. To characterize the conditions at the area-level, values were derived from the mean of two vertical profiles taken in each area/bay (one nearshore and one towards the central parts of the area) during each sampling event (Figure S1).

### 3.6 Species identification and biometrics

In the laboratory, ethanol-preserved mysids were identified to species level based on morphological characteristics following Köhn et al. (1992). All intact specimens were digitized using an EPSON Perfection V600 flatbed scanner (Seiko Epson Corp., Suwa, Japan). Total body length was measured from the tip of the rostrum to the base of the telson using the segmented line tool in ImageJ (Schneider, Rasband and Eliceiri 2012), adhering to standard operating procedures for mysid analysis (US EPA 2015)

Ethanol preserved *G. aculeatus* caught in the reference benthic traps were measured to the nearest millimetre (total length) using a measuring board. These measurements were used to evaluate potential physical constraints on trap entry. Since the light traps utilized fixed entrance slots (5-7 mm), monitoring the size structure of the ambient stickleback population allowed us to determine whether the seasonal decline in light trap catches was a result of biological factors (e.g., reduced phototaxis) or a methodological artifact caused by fish growing too large to pass through the entrance slits.

### 3.7 Size distribution of mysids

Population structure was visualized using length-frequency histograms faceted by sampling date. The primary aim of this analysis was to evaluate the possible size selectivity of the light traps. By examining the length distributions across the season, we assessed whether the gear successfully captured the full ontogenetic range of the target species, from newly released juveniles to large overwintering adults. This validation was essential to confirm that the traps function as an unbiased monitoring tool capable of tracking the entire life cycle.

### 3.8 Statistical analysis

#### 3.8.1 General procedures

All statistical analyses and data visualizations were performed using R statistical software (v. 4.5.0; R R Core Team 2022). Data wrangling and visualization were conducted using the tidyverse suite of packages (Wickham *et al*. 2019). R-scripts and raw data for the analyses are provided in the Supplementary Information (SI_file_package_2.zip).

#### 3.8.2 Exploratory analysis and variable selection

Prior to modelling, we assessed collinearity among environmental variables (*Water temperature, Turbidity*, *Night duration*, and *Julian day*) by examining pairwise correlation matrices. We identified strong collinearity between *Night duration* and *Water temperature* (Pearson’s r > 0.8, Figure S2) and between *Night duration* and *Julian day.* Including these correlated variables simultaneously resulted in Variance Inflation Factors (VIF) > 6, necessitating a selection protocol to identify the primary mechanistic driver of catch rates.

To resolve this, we adopted a mechanism-based selection protocol. For light traps, where catch rates could theoretically be driven by either metabolic activity (*Temperature*, (Clarke and Johnston 1999; Gillooly *et al*. 2001; Stehfest, Lyle and Semmens 2015)) or the visibility of the attraction stimulus (*Night duration*), we performed a comparative model selection analysis. Single-predictor models were ranked using Akaike Information Criterion (AIC). For all taxa in light traps, models based on night duration were statistically superior (ΔAIC > 6, Table S1), confirming that the seasonal catch pattern was primarily driven by the duration of the light attraction window. Conversely, for the benthic trap analysis, water temperature was selected à priori as the explanatory variable. Since these unlit, passive gears rely entirely on the random movement of organisms, and night duration has no mechanistic function in attracting fish to unlit traps, we interpreted the seasonal trend according to metabolic activity rather than performing a redundant statistical comparison against a non-mechanistic proxy. All continuous predictors were mean-centred to facilitate model convergence.

#### 3.8.3 Modelling framework and validation

To analyse the catch data, we fitted Negative Binomial Generalized Linear Mixed Models (GLMMs) with nested random effects (*Station* within *Area*) via the *glmmTMB* package (Brooks *et al*. 2017). Preliminary interaction analyses indicated that the two mysid species exhibited distinct temporal dynamics and required separate models. Conversely, the two stickleback species (*G. aculeatus* and *P. pungitius*) showed similar responses and were pooled for light trap analyses.

We restricted benthic trap models to the dominant *G. aculeatus* due to low frequency of occurrence and zero-inflation in the *P. pungitius* catch (17.7 %, Table S4).

To evaluate the effect of stickleback presence on mysid catch, we modelled the catch rates of the numerically dominant mysid (*N. integer*) as a function of pooled stickleback abundance. To determine if predator effects were specific to sticklebacks or a general consequence of high fish densities, we fitted an identical control model using the abundance of sympatric pipefish (*Nerophis ophidion* and *Syngnathus typhle*).

We validated model assumptions, including overdispersion and spatial autocorrelation, using the *DHARMa* package (Hartig 2022). No significant violations were detected that necessitated altering the parsimonious random effect structure. Comprehensive details regarding model selection, species pooling criteria, spatial autocorrelation tests, and full model equations (Eq. S1–S3) are provided in Supplementary Information section S4.

#### 3.8.4 Sampling optimization and power analysis

To determine the optimal sampling effort required to monitor littoral communities, we employed a two-stage simulation approach based on parameters extracted from our empirical catch data. We selected *N. integer* as the primary model species for optimizing effort due to its high abundance and aggregation, while also evaluating pooled stickleback abundance (*G. aculeatus* and *P. pungitius*) as a methodological benchmark to assess gear performance across different dispersion profiles. First, we applied the precision optimization method of Hewitt *et al*. (1993) to optimize the cost-precision trade-off, identifying the asymptote where further increases in sample size yielded negligible reductions in relative standard error (CV).

Second, utilizing the optimal effort identified (n = 10 traps per area), we performed a hierarchical power analysis using a discrete “step-change” approach (Hawley *et al*. 2019) suitable for short-lived invertebrates and mesopredatory fish. We simulated varying magnitudes of population declines (50% to 90%) and recruitment increases (1.5-fold to 5-fold) to assess the monitoring design’s detection sensitivity while accounting for spatial variability. Full simulation procedure, variance parameterization, and Generalized Linear Mixed Model (GLMM) equations are detailed in Supplementary Information section S5.

## 4 Results

### 4.1 Environmental conditions

Surface water temperatures followed a typical seasonal progression, rising steadily from approximately 5°C in late April to >20°C by early July (Figure S1). This warming coincided with a gradual decline in dissolved oxygen concentrations, which decreased from ∼14.5 mg/L at the start of the study to ∼9.5 mg/L in July. Salinity remained stable (∼6.2 PSU) across most sampling areas throughout the study period, with the exception of a transient freshwater pulse recorded in Area B during late May. This was caused by an artificial dam rupturing in a local stream. Chlorophyll-a fluorescence and turbidity generally remained low, though sporadic peaks were observed in late June, indicative of localized plankton blooms or resuspension events.

### 4.2 Catch composition

Apart from mysids and sticklebacks, the light traps also captured high numbers of straight-nosed pipefish (*Nerophis ophidion*), yielding a total abundance comparable to that of *G. aculeatus* (Table S6). Due to their slender body morphology (typically 3–5 mm in diameter), these fish were not effectively excluded by the netting and occurred in both trap types at similar proportions. Similarly, although the netting significantly reduced stickleback entry, a small number of individuals managed to penetrate minute gaps in the mesh, meaning that the modified traps were not completely stickleback-free. Other bycatch species in the unmodified traps included broad-nosed pipefish (*Syngnathus typhle*), sand goby (*Pomatoschistus minutus*), two-spotted goby (*Gobiusculus flavescens*), viviparous eelpout (*Zoarces viviparus*), rock gunnel (*Pholis gunnellus*), and European perch (*Perca fluviatilis*) and a few unidentified fish larvae (Table S6).

### 4.3 Effects of trap design and stickleback pooling

We first assessed whether stickleback species responded differently to environmental conditions using a model that included species-interaction terms. The interaction between species identity and night duration was non-significant (p = 0.32, Table S3), as was the interaction with temperature (p = 0.69, Table S3). These results indicate that *G. aculeatus* and *P. pungitius* exhibit statistically indistinguishable responses to light trap stimuli. Consequently, abundances of both species were pooled for the subsequent analysis of drivers and statistical power.

Trap design significantly influenced these pooled catch rates. Unmodified traps, which allowed stickleback entry, caught approximately 5 times more sticklebacks than modified traps equipped with exclusion netting (IRR = 5.37 (e^1.68^), p < 0.001, Table 1). However, the presence of these mesopredators in unmodified traps significantly reduced the catch efficiency of mysids. Allowing stickleback entry reduced mysid catches by approximately 85% for *Neomysis integer* (IRR = 0.15, p = 0.001; Table 2, Figure 4A) and 82% for *Praunus flexuosus* (IRR = 0.18, p < 0.001, Table 2, Figure 5A). The negative impact of mesopredator density was further confirmed by the “Mod1b” model (Table 2), which showed a steep decline in *N. integer* catches as stickleback abundance increased within the traps (Figure 4C). In stark contrast, we found no evidence of a similar depletion effect using pipefish (*Nerophis ophidion* and *Syngnathus typhle*). Despite being the numerically dominant fish taxa in the traps (Table S6), pipefish abundance was not a significant predictor of mysid catch rates (GLMM: β= -0.006 ± 0.02 SE, z = --0.26, p = 0.80).

**Figure 4.**
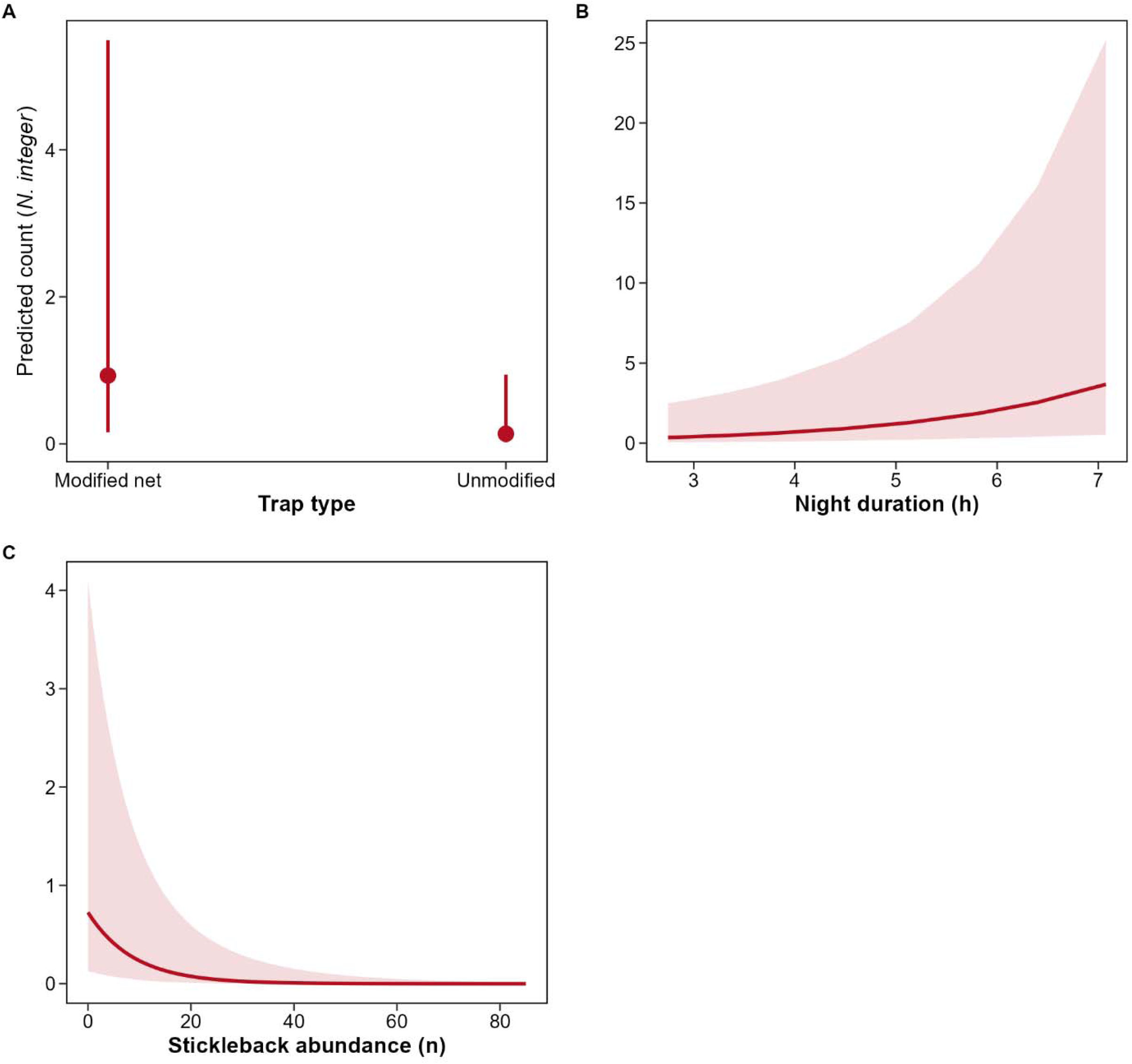
Marginal means effect plots from the N. integer model (Table 2, Mod1). The plot shows the predicted catch of N. integer in the modified and unmodified light trap (A) and the effect of night duration in the modified light traps (B). The effect of stickleback abundance on the catch of N. integer at mean night duration (C) is derived from the model where Trap type has been replaced by stickleback abundance (Table 2, Mod1b).

**Figure 5.**
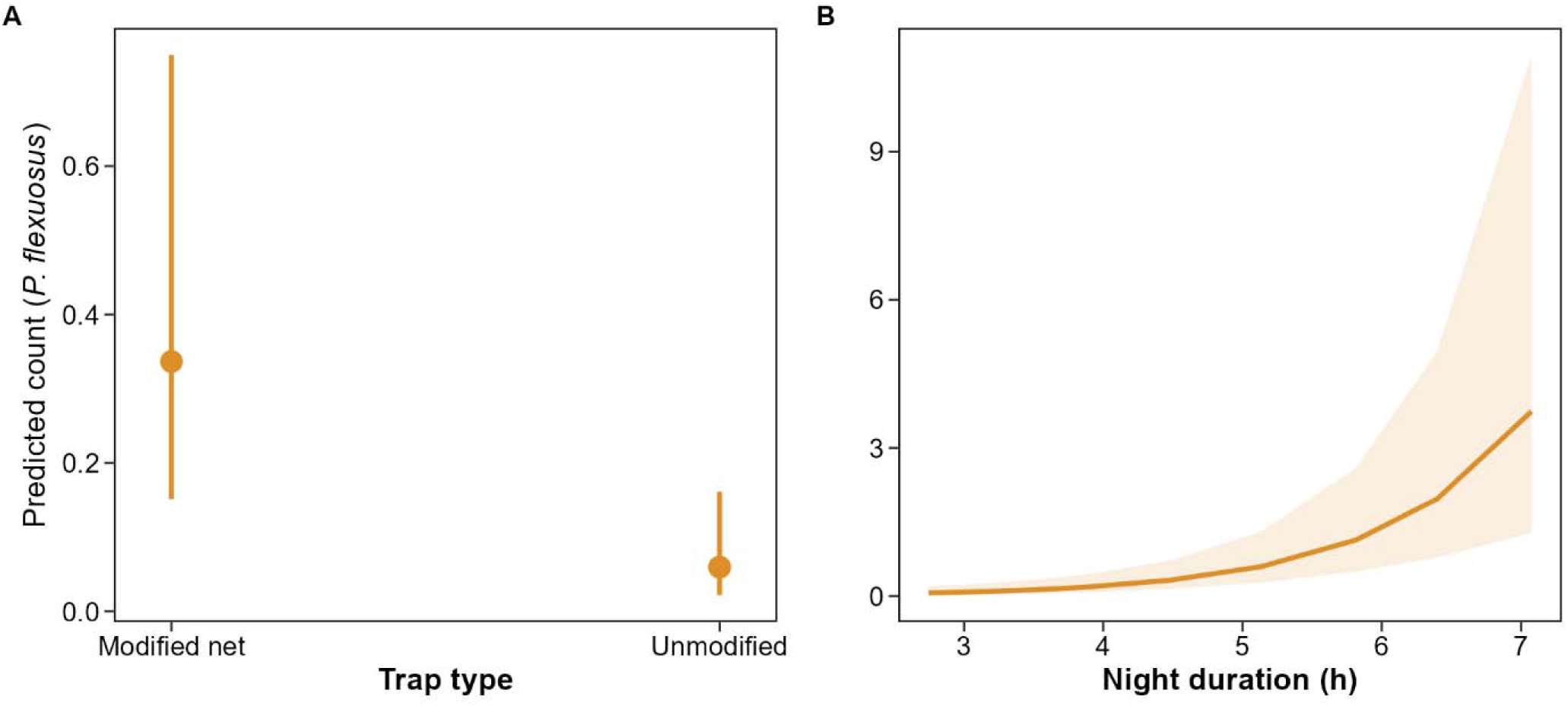
Marginal means effect plots from the P. flexuosus model (Table 2, Mod2). The plot is showing the predicted catch of N. integer in the modified and unmodified light trap (A) and the effect of night duration in the modified light traps (B).

**Table 1.**
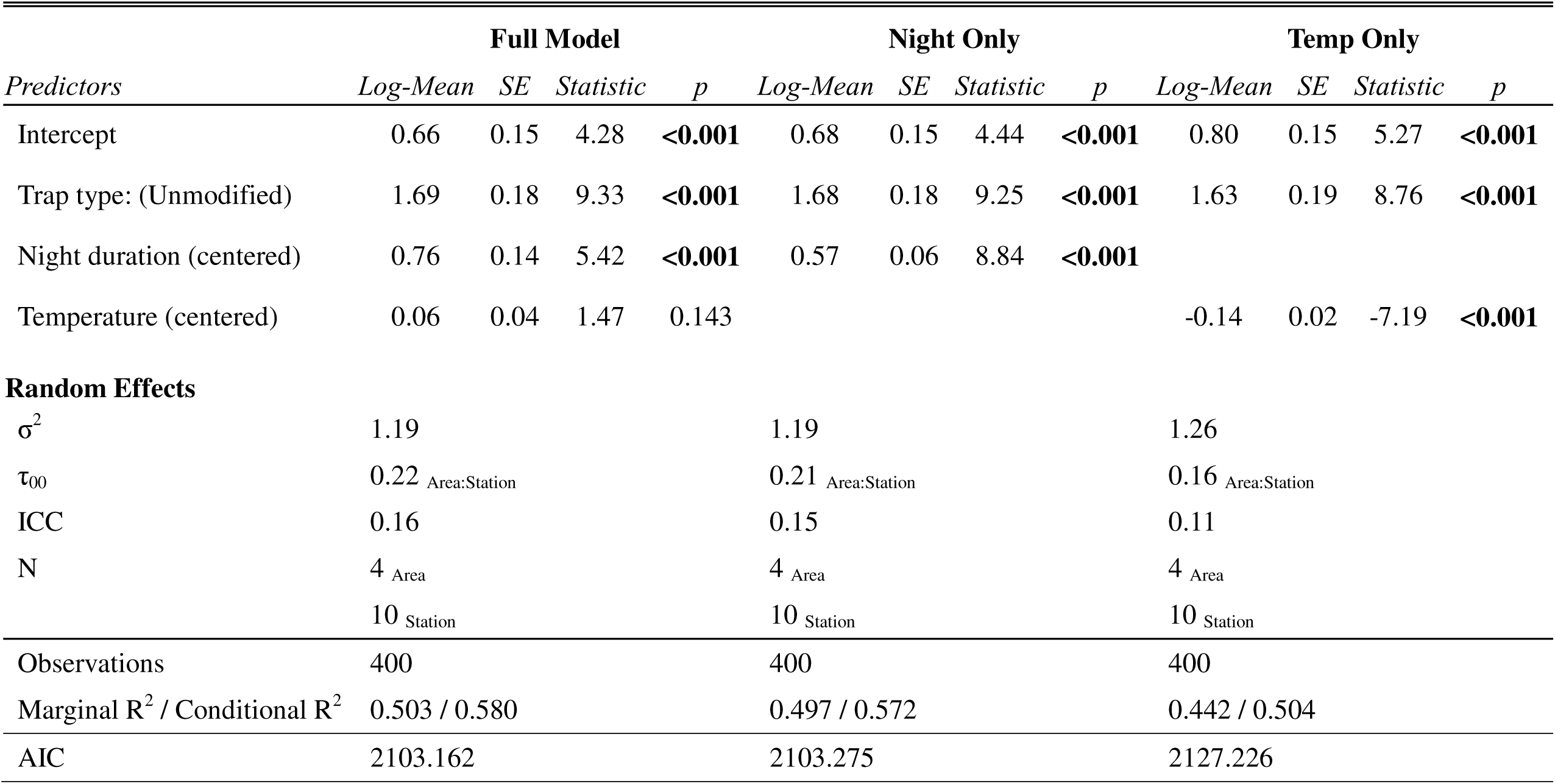
Model selection results for stickleback abundance (species pooled) in light traps. Three competing models were fitted to disentangle the effects of night duration and water temperature, which were strongly collinear. “Full Model” includes both predictors; “Night Only” includes only night duration; “Temp Only” includes only temperature. Akaike Information Criterion (AIC) indicates the “Night Only” model is the most parsimonious (lowest AIC). Significant values are outlined in bold.

**Table 2.** Summary of generalized linear mixed model (GLMM) results estimating the abundance of Neomysis integer and Praunus flexuosus in response to trap type (Mod1 and Mod2) and for N. integer, stickleback abundance (Mod1b). Standard errors are given within parentheses, and significant effects are indicated by bold text.

|  | Neomysis (Mod1) |  |  | Praunus (Mod2) |  |  | Neomysis (Mod1b) |  |  |
| --- | --- | --- | --- | --- | --- | --- | --- | --- | --- |
| Predictors | <sup>a</sup>IRR | Statistic | p | <sup>a</sup>IRR | Statistic | p | <sup>a</sup>IRR | Statistic | p |
| Intercept | 0.93<br>(0.84) | -0.08 | 0.934 | 0.34<br>(0.14) | -2.67 | **0.008** | 0.28<br>(0.25) | -1.14 | 0.161 |
| Trap type:<br>(Unmodified) | 0.15<br>(0.09) | -3.20 | **0.001** | 0.18<br>(0.08) | -3.74 | **<0.001** |  |  |  |
| Night duration (c) | 1.72<br>(0.32) | 2.89 | **0.004** | 2.58<br>(0.48) | 5.13 | **<0.001** | 2.26<br>(0.49) | 3.77 | **<0.001** |
| Stickleback<br>abundance (c) |  |  |  |  |  |  | 0.89<br>(0.03) | -3.49 | **<0.001** |
| $\sigma^2$ | | 2.49 | | | 2.01 | | | 2.45 | |
| $\tau_{00}$ | | 2.80 Station:Area | | | 2.71 Station:Area | | | 3.02 Station:Area | |
|  |  | 2.39 Area |  |  | 0.00 Area |  |  | 2.26 Area |  |
| ICC |  | 0.68 |  |  | 0.57 |  |  | 0.68 |  |
| N |  | 10 Station |  |  | 10 Station |  |  | 10 Station |  |
|  |  | 4 Area |  |  | 4 Area |  |  | 4 Area |  |
| Observations |  | 400 |  |  | 400 |  |  | 400 |  |
| Marginal R<sup>2</sup> /<br>Conditional R<sup>2</sup> |  | 0.165 / 0.729 |  |  | 0.355 / 0.725 |  |  | 0.263 / 0.766 |  |

<sup>a</sup>IRR = Incidence Rate Ratio; (c) = Mean centered variable

### 4.4 Environmental drivers of mysid and pooled stickleback catch rates

Model selection confirmed that night duration was the primary driver of catch rates for both mysid species in light traps, outperforming water temperature (Table S1). For *N. integer*, catch rates were positively associated with night duration in models including *Trap type* and *Stickleback abundance* as covariates respectively (IRR = 1.7, p = 0.004 and IRR = 2.3, p <0.001, Table 2), indicating that abundance in traps increased as the dark “attraction window” lengthened (Figure 4B). *P. flexuosus* exhibited an even stronger sensitivity to light conditions. The effect of night duration was highly significant (p < 0.001), with the incidence rate ratio indicating a 2.6-fold increase in catch for every unit increase in centred night duration (Table 2; Figure 5B). For the pooled stickleback metric, model selection favoured the “Night-Only” model (ΔAIC > 23.9 against the temperature model; Table 1, Table S1), confirming that the seasonal catch pattern was driven by the night duration rather than temperature. This model estimated a strong positive relationship between night duration and catch (β = 0.57, p < 0.001, Table 1).

### 4.5 Disentangling methodological constraints from biological activity in *G. aculeatus*

To distinguish between changes in gear efficiency and changes in actual population density, we compared the environmental drivers of *G. aculeatus* catches in unmodified light traps against those in benthic traps. Since benthic traps are passive gears compared to the light traps which actively attract organisms, they served as a reference to verify whether the summer decline in sticklebacks observed in light traps represented a true population decline or just a decreased catchability of the light traps

In the light traps, catch rates were decoupled from the expectation of increasing mobility over the sampling season. Despite rising water temperatures (which typically increase fish activity) catches peaked in early spring and declined as the season progressed. Model selection favored *Night duration* over *Water temperature* as the primary driver (Table 1), confirming that catches were regulated by the duration of the dark “attraction window” rather than biological abundance.

In contrast, benthic traps revealed the inverse pattern. Catch rates in these passive gears showed a strong positive response to *Water temperature* (IRR = 1.18, p < 0.001; Table S8; Figure 6A), consistent with the expected increase in ectothermic activity during the warmer summer months. While night duration was highly correlated with benthic catches, the relationship was negative (catches decreased with increasing night duration; Figure 6B), directly opposing the light trap trend. This divergence confirms that the summer decline in light trap catches was a consequence of shortening nights, while the benthic traps correctly tracked the high biological activity of the warm season.

**Figure 6.**
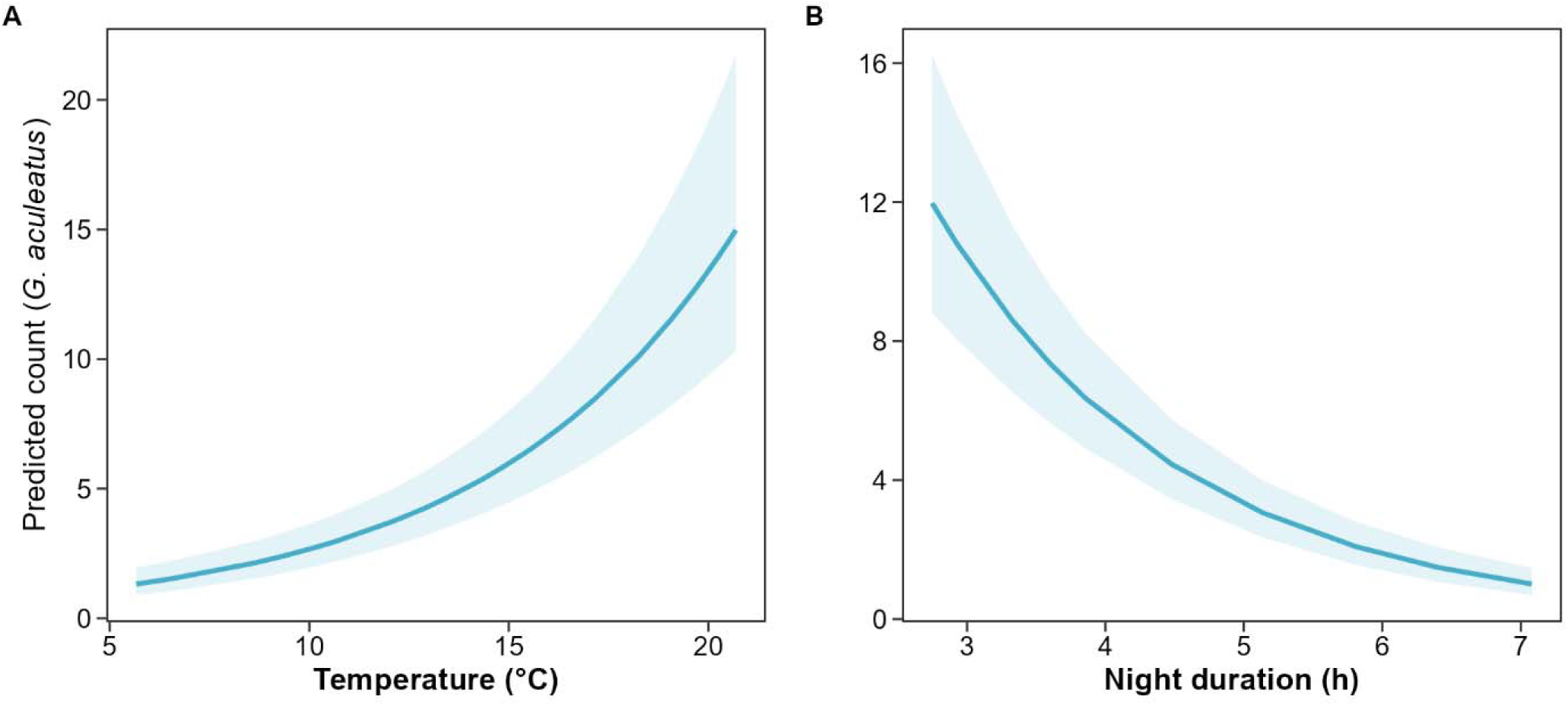
Marginal means effect plots from models predicting G. aculeatus catch in benthic traps. Panel A shows the effect of water temperature (Table S 8, “Temp Only model”) and panel B the effect of night duration.

### 4.6 Size selectivity and population structure

Length-frequency analysis confirmed that the light traps successfully sampled the entire life cycle of the target mysid species, ruling out size-based exclusion as a cause for seasonal declines (Figure 7). *Neomysis integer* populations were dominated by large overwintering adults (mean length ∼11–13 mm) in April and May. A distinct recruitment pulse of juveniles (<8 mm) appeared in mid-June and dominated the population structure into July. Catches of *P. flexuosus* were restricted almost exclusively to the spring period, peaking in mid-May with large, mature individuals (mean length ∼16 mm). Coincident with the onset of the shortest nights in June, *P. flexuosus* virtually disappeared from the catch, dropping to negligible levels (≤ 1 individual per sampling occasion), despite the continued presence of *N. integer*.

**Figure 7.**
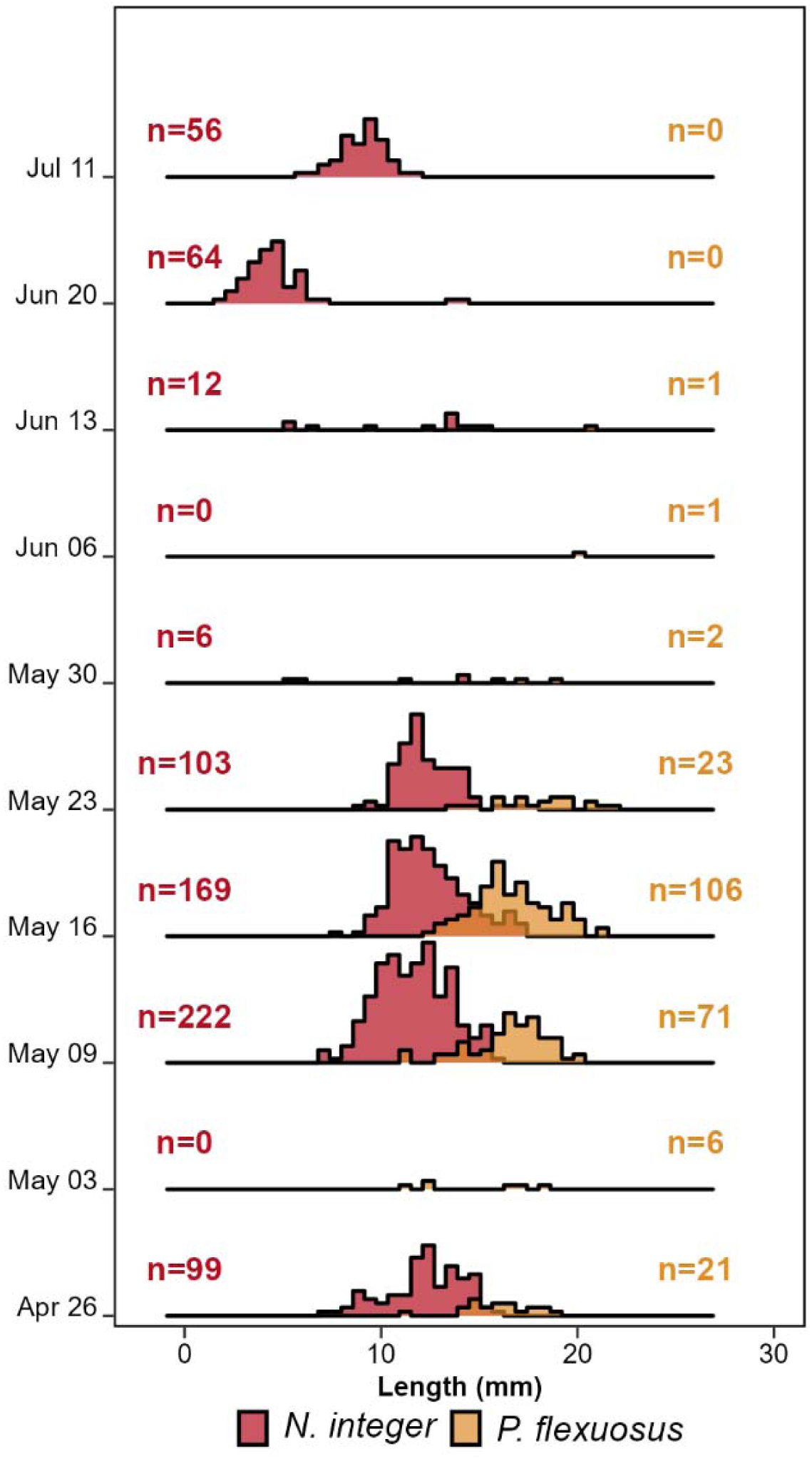
Seasonal size-frequency distributions of the dominant mysid species, Neomysis integer (red) and Praunus flexuosus (yellow), caught in modified light traps from April to July. Histograms are normalized to show relative size structure within each sampling date (i.e., each distribution scales to a total proportion of 1). Annotations (n) indicate the total raw abundance (number of individuals) caught for each species per date.

### 4.7 Sampling optimization and power analysis

Analysis of sampling precision (Figure S4) revealed a clear distinction in monitoring performance between the two functional groups. For the highly aggregated *N. integer*, the coefficient of variation (CV) was initially high but declined steeply with increasing effort, identifying *n* = 10 traps as the optimal cost-precision-trade-off, where further increases in sampling effort yielded only marginal gains in precision.

In contrast, the pooled stickleback guild exhibited significantly higher baseline precision due to their more uniform distribution (θ = 0.62) and higher absolute abundance. The CV stabilized rapidly, reaching robust levels (CV < 1) by 6 traps. Consequently, applying the mysid-optimized effort (*n* = 10) to the stickleback guild represents a conservative sampling intensity. While this effort merely optimized cost-efficiency for the patchy mysids, it provided a safety margin for the sticklebacks, yielding precision that exceeded the stabilization threshold for this group.

Power analysis for the highly aggregated mysid *N. integer* demonstrated low statistical power (< 0.25) to detect moderate declines of 30–50% across all simulated sampling efforts (5–20 traps), with no appreciable gain from increasing effort (Figure 8A). Reliable detection (> 0.80 power) for mysids was restricted only to substantial population declines of > 90%.

**Figure 8.**
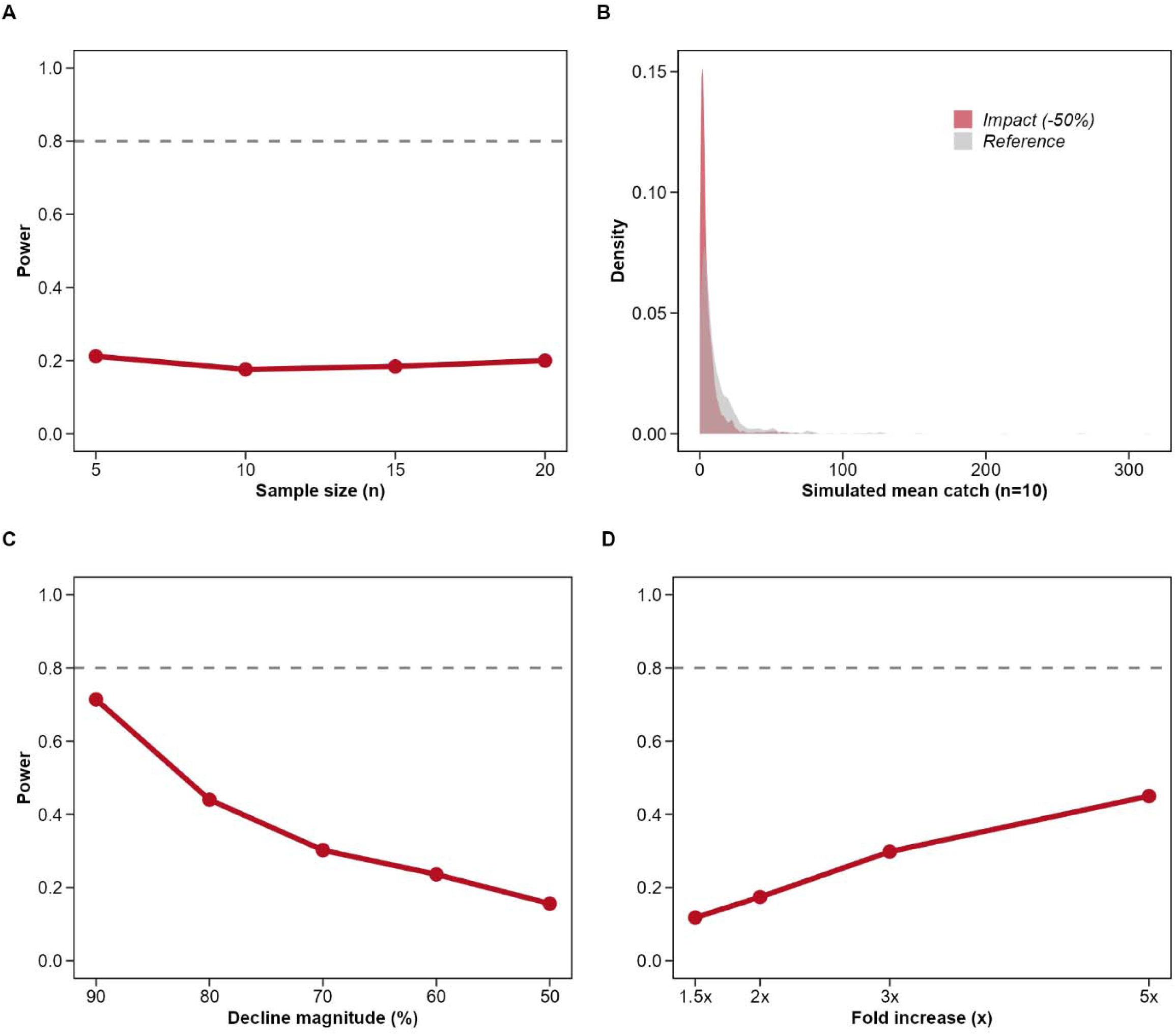
Statistical power analysis for detecting abundance changes in Neomysis integer using Generalized Linear Mixed Models (GLMMs). (A) Power to detect a 50% population decline relative to spring baseline ( µ = 5 ind./trap) across increasing sampling effort. (B) Simulation density plot based on 10 traps, showing the distribution of mean catches for Reference (grey) versus Impact (red) scenarios. (C) Sensitivity analysis for detecting declines of 10–50% at a fixed effort of n=10. (D) Power to detect population increases (1.5 to 5-fold) at n=10.

The high degree of overlap in sampling distributions of the “Reference” and “Impact” scenarios demonstrated that this low power is driven by the high natural dispersion (θ) which effectively masks the signal to detect population size changes (Figure 8B). Consequently, reliable detection of mysid declines was restricted to substantial collapse scenarios; power only exceeded 0.60 when the population declined by ≥ 90% (Figure 8C). Similarly, population increases were difficult to confirm, with even a 5-fold increase in abundance yielding detection power of only ∼0.45 (Figure 8D).

In contrast, the monitoring design proved highly robust for the sticklebacks, despite being simulated at the same low starting density (µ = 5) as the mysids. Power increased linearly with effort, crossing the 0.80 threshold at n = 16 traps (Figure 9A). At the chosen effort of n=10, the design achieved moderate power (0.62) to detect a 50% decline which also was supported by a distinct separation between the reference and impact sampling distributions (Figure 9B) The method was highly sensitive to larger changes in the fish community, detecting declines of 70% and 3-fold population increases with > 0.90 power (Figure 9C, D).

**Figure 9.**
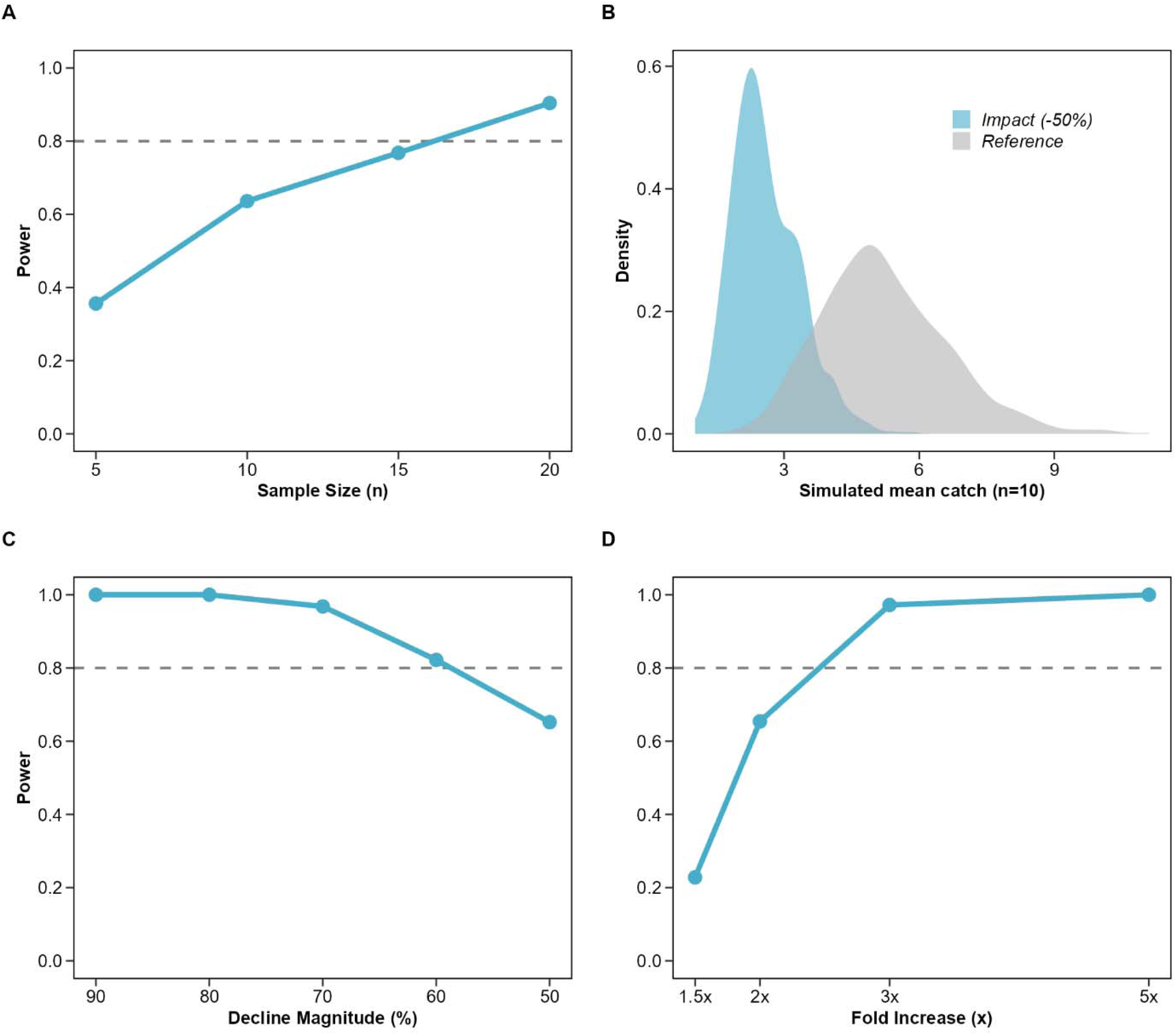
Statistical power analysis for detecting abundance changes in sticklebacks (G. aculeatus + P. pungitius) using Generalized Linear Mixed Models (GLMMs). (A) Power to detect a 50% population decline relative to spring baseline ( µ = 5 ind./trap) across increasing sampling effort. (B) Simulation density plot based on 10 traps, showing the distribution of mean catches for Reference (grey) versus Impact (red) scenarios. (C) Sensitivity analysis for detecting declines of 10–50% at a fixed effort of n=10. (D) Power to detect population increases (1.5 to 5-fold) at n=10.

## 5 Discussion

### 5.1 Light availability drives light trap efficiency

Our results demonstrate that light traps are effective in capturing both littoral mysids and mesopredatory fish, like sticklebacks, but the efficacy of the traps in temperate shallow waters is regulated primarily by the seasonal availability of darkness. The strongest evidence for this is the divergence in drivers between light-based and passive gears. While unlit benthic traps tracked the expected increase in catch rates over the season, due to biological increases in fish activity during warmer months (Jacobsen *et al*. 2002; Linløkken and Haugen 2006), light traps exhibited the opposite pattern.

The peak in light trap catches during the darkest months and the distinct decline during the spring/summer confirms that the method is constrained by contrast reduction. As night duration shrinks below critical thresholds, the operational window becomes too short to attract organisms effectively. This finding aligns with Lindquist & Shaw (2005), who reported that environmental factors attenuating light transmission (e.g., turbidity) or reducing contrast (e.g., background illumination) significantly reduce Catch Per Unit Effort (CPUE) for larval fishes. Similarly, Meekan *et al*. (2001) established that ambient light levels (e.g., lunar cycles) critically modulate the effective attraction radius of light traps. Our data extends this by showing that in high-latitude summers, the ambient light from short nights effectively mimics the lunar inhibition effect described in other geographical locations (Verheijen 1960; Hickford and Schiel 1999).

### 5.2 Biotic interactions and gear selection

We observed a strong negative relationship between pooled stickleback abundance and mysid catches. This reduction is likely driven by a behavioural avoidance and possibly also “in-trap” predation. As sticklebacks are voracious visual predators, predation in the traps cannot be excluded. However, field observations from Danish hypertrophic lakes suggest that adult *G. aculeatus* primarily consume juvenile *N. integer* (3–4 mm) rather than the larger size classes caught in our modified traps (Søndergaard, Jeppesen and Aaser 2000).

This discrepancy between stickleback prey-size selectivity and the actual size distribution of mysids in our samples suggests that direct predation is unlikely to explain the magnitude of the decline. In laboratory experiments, Lindén *et al*. (2003) observed that *N. integer* reduce their swimming activity and *P. flexuosus* seek shelter in vegetation in the presence of perch (*P. fluviatilis*). Both mysid species also reduced their food ingestion but only when the chemical cues were coupled with visual stimuli. The fact that our unmodified traps, which allowed stickleback entry, caught ∼85% fewer mysids than exclusion traps suggests that this avoidance-effect is operating within the gear itself. This is a sampling bias frequently observed in trap-based monitoring where predation and behavioural avoidance distorts abundance indices (Bacheler 2024 and references therein)

The species-specific nature of this biotic interference is further supported by our findings regarding the lack of association between pipefish and mysid catches. Although the pipefish were as abundant as sticklebacks and are specialized suction feeders capable of capturing elusive prey (Van Wassenbergh *et al*. 2007; Sundin *et al*. 2011), their presence had no detectable impact on mysid catch rates. This discrepancy likely reflects differences in predator recognition and foraging behaviour. Sticklebacks are active, visual hunters that likely trigger chemically or visually mediated escape responses in mysids. Conversely, the slow-moving pipefish appears to not elicit the same avoidance response as the more active sticklebacks. Furthermore, the relatively small gape size of the pipefish likely restricts them from consuming the adult mysids that dominated our catch during most of the study period, effectively ruling out in-trap predation as a source of bias for this species.

### 5.3 Implications for monitoring and statistical power

Ideally, the capture efficiency of a new sampling method would be validated against a known absolute abundance or a standard reference gear (ground truthing). Previous studies have successfully cross-validated light traps against vertical net tows conducted adjacent to docks and piers (Brown *et al*. 2017). However, while net validation is feasible in accessible water where avoidance behaviour is assumed to be constant, this assumption likely fails in structurally complex littoral zones. The variable density of vegetation and irregular rocky substrates like in the Baltic archipelagos physically impede active gears, causing filtration efficiency to fluctuate unpredictably. Furthermore, even where comparisons are possible, the relationship between light trap catch and net density is often non-linear (Brown *et al*. 2017), suggesting density-dependent catchability. By contrast, light traps function as passive, standardized samplers. When corrected for predator exclusion and night duration in statistical models, they offer a consistent CPUE index in complex habitats where active trawling is operationally inconsistent. Consequently, rather than calibrating against a biased reference method, we evaluated the utility of light traps based on their internal consistency and statistical power to detect relative changes in abundance

Our power simulations highlight that light traps seem to be ineffective as a monitoring method for littoral mysids due to their patchiness. For the highly aggregated mysid *N. integer*, the method demonstrated low statistical power (< 0.25) to detect even moderate population declines of 50%. This has implications for management as it indicates that abundance-based monitoring for swarming aquatic invertebrates is constrained by the high overlap in sampling distributions (Figure 8B). Specifically, the probability distribution of a population that has declined by 50% overlaps almost completely with the natural variability of the population, rendering the two states statistically indistinguishable. This suggests that light traps cannot robustly assess the status of mysid populations between the start and end of a standard 6-year MSFD assessment cycle (European Commission 2008), where cumulative declines of this magnitude might be expected. Instead, for these species, the gear functions strictly as an “ecosystem-shift” indicator capable of reliably detecting only significant population declines or increases.

In contrast, the comparative analysis with sticklebacks validates the light trap as a precise tool for less aggregated taxa. For these, currently abundant mesopredators, the method achieved robust power (> 0.80) to detect a 60% difference at simulated low abundances. This confirms that the gear is sufficiently precise to detect declines of this magnitude even when at low population densities. Consequently, while light traps yield high-resolution data for sticklebacks, monitoring programs targeting mysids using light traps must accept that statistical significance will likely be limited to distinguishing between “present” and “functionally extinct”, rather than tracking subtle trends.

However, it should be recognized that power analyses focused on single parameters like abundance do not capture the full scope of ecological information gathered during a continuous monitoring program (Lindenmayer and Likens 2010). Integrated metrics, such as shifts in size distribution, timing of recruitment pulses, and seasonal phenology interact to paint a coherent picture of organismal status that simple count data may miss. Hence, poor statistical power for abundance alone does not necessarily disqualify a sampling method. As debated widely in the literature (Mapstone 1995; Seavy and Reynolds 2007), the utility of power analysis in environmental monitoring is not absolute; the framework should be used to contextualize data limitations rather than to strictly define methodological validity.

### 5.4 Light traps as a cold-season alternative for stickleback monitoring

An unexpected but valuable finding was the superior performance of light traps for monitoring sticklebacks during the cold season. In early spring, when water temperatures were low, benthic trap catches were negligible. In contrast, light traps yielded their highest catches during this period. This suggests that the visual stimulus is sufficient to overcome the reduced activity of fish in cold water. At low temperatures, fish swimming performance and reactive distances typically decrease, rendering passive gears that rely on voluntary movement ineffective (Bacheler and Shertzer 2020). Our results indicate that light attraction can override this thermal inhibition, filling a critical methodological gap for winter/spring monitoring when passive gears underperform. Since sticklebacks have been implicated in predation on early life-stages of e.g. northern pike (Bergström *et al*. 2015; Donadi *et al*. 2020) that spawn in early spring, light traps can be particularly useful for studying trophic and behavioural interactions between these species during a time and space where these interactions are anticipated to occur.

### 5.5 Conclusions

Our study evaluated light traps as a tool for monitoring mysids, though our results demonstrate that they are significantly more effective for monitoring mesopredatory sticklebacks. While the traps provide a unique capacity to collect mysid data that is currently unavailable, their primary strength lies in the precise monitoring of sticklebacks during the dark winter and spring periods in temperate regions. We recommend the use of light traps for these species, provided that abundance indices are modelled to account for night duration. Furthermore, the use of physical exclusion barriers is essential when specifically attempting to quantify mysid populations.

## Supporting information

5D design files for the quatrefoil light trap and custom light source

## 6 Acknowledgements

We would like to thank Johan Wenngren (Askö, Sweden) and associates for carrying out the sampling and assistance in modifying the light traps; the Stockholm University Baltic Sea Centre, Askö laboratory for providing housing and equipment, Douglas Jones, NIRAS AB, Sweden and Björn Rogell, Swedish University of Agricultural Sciences for valuable comments on the manuscript.

During the preparation of this manuscript, the author used Google Gemini 3.0 to improve the readability and flow of the text, as well as to assist with troubleshooting R scripts and refining statistical interpretations. After using this tool, the author rigorously reviewed, tested, and edited all generated text and code as needed. The author takes full responsibility for the ultimate content, analyses, and conclusions presented in this publication.

## 7 Author contributions

Conceptualization: M.O; Methodology: M.O; Data curation: M.O; Formal analysis: M.O; Funding acquisition: M.O, Project administration: M.O; Visualization: M.O; Writing – original draft: M.O; Writing – review & editing: M.O.

## 8 Ethical statement and wildlife welfare

All applicable international, national, and/or institutional guidelines for the care and use of animals were followed. The fish sampled and handled in this study complied with the standards and procedures stipulated by the Swedish Ministry of Agriculture and the Swedish Board of Agriculture. The research was conducted under a formal permit for animal experimentation, which was approved by the Linköping Regional Animal Ethics Committee (DNR: 23379-2022). All measures possible were taken to follow the 3R tenets (Replacement, Reduction, and Refinement) as part of the ethical vetting process for our permit application. Specifically, refinement was achieved through the use of passive, non-lethal gear (light traps and fish traps). The study followed the taxon-specific guidelines for the ethical treatment of fishes as outlined by the Stockholm University Baltic Sea Centre (document number: SU-484-0009-22). Following identification and quantification, all bycatch was monitored for normal swimming behaviour and released immediately at the site of capture.

