## Supporting information for "Optimizing light traps for littoral mysids and mesopredatory fish in the Baltic Sea: environmental drivers and seasonal monitoring efficacy"

Martin Ogonowski*

Department of Aquatic Resources, Institute of Freshwater Research, Swedish University of Agricultural Sciences, Stångholmsvägen 2, 178 93 Drottningholm, Sweden

#### S1. Light trap design and construction

The quatrefoil light traps were custom-manufactured at the Department of Geological Sciences, Stockholm University. The main body was constructed from CNC-machined 10 mm polycarbonate (PC) sheeting and tubing (outer diameter = 75 mm). To ensure positive buoyancy and a vertical orientation in the water column, a 50 mm layer of extruded polystyrene foam was adhered to the trap's upper surface.

The collection mechanism consisted of a stainless-steel vessel (formerly a commercial baking pan; Homikit) secured to the polycarbonate body via custom PC clamps. To facilitate drainage upon retrieval, a 10 mm hole was drilled into the side of the collection vessel and covered with a welded 0.5 mm stainless steel mesh.

#### S2. Light trap illumination

We used light-emitting-diodes (LED) instead of the commonly used chemical light sticks to achieve a more consistent illumination profile (Adams *et al.* 2023). The light source consisted of a 3D-printed white PLA tube holding four green LEDs, powered by a 600 mAh rechargeable Li-ion battery (Bonai, Shenzhen Tuoman Technology Co., Ltd., Shenzhen, China). This assembly was housed within a waterproof, polypropylene screw-cap container (70 mL, 55 × 44 mm; Sarstedt AG, Nümbrecht, Germany). To ensure water resistance and create a uniform, diffuse light field, the electrical connections and LEDs were encased in translucent silicone. Complete 3D-design files for the light trap body and light source assembly are provided in the Supplementary Material (SI_file_package_1.zip).

#### S3. Spectral characteristics of mysids and illumination intensity

The LEDs were selected for a peak emission (λ*_peak_* of 525 nm, targeting the spectral sensitivity maxima (*S*(λ)*_max_*) of *Neomysis integer* (525–535 nm) and *Praunus flexuosus* (505–515 nm) (Lindström 1992).

Light intensity was characterized in a dark room using a photometer (Model UT383; Uni-Trend Technology, Dongguan, China). Due to the low-intensity design, illuminance (*E_v_*) was measured at a fixed distance of 10 cm (8 lux). To report biologically relevant light levels, standard lux values were converted to "mylux" following Boscarino et al. (2009). The conversion factor was derived from the ratio of mysid sensitivity (*S_mysid_* ≈ 1.0) to human photopic sensitivity (*V_λ_* ≈ 0.79) at 525 nm. Consequently, the measured illuminance was multiplied by a factor of 1.27. This resulted in a calculated intensity of **~**0.10 mylux at 1 m (derived from an extrapolated value of 0.08 lux). This intensity approximates natural moonlight levels (Kyba, Mohar and Posch 2017), likely minimizing negative phototaxis.

#### S4 Statistical modelling framework, species grouping, and validation

We employed a stepwise modelling approach to determine species groupings. First, to determine if *N.integer* and *P. flexuosus* could be analysed as a single functional group, we fitted a combined model including interaction terms for *Species* × *Trap type* and *Species* × *Night duration* (Table S2, Eq S1). As described in Table S2, significant interactions necessitated the use of separate models for each mysid species to accurately capture distinct temporal and behavioural dynamics.

Similarly, we assessed whether the two stickleback species (*G. aculeatus* and *P. pungitius*) exhibited distinct responses to environmental conditions in light traps. We fitted a preliminary GLMM including species-interaction terms (*Species* × *Night duration*; Species × *Temperature,* Table S3). Finding no significant interactions (p > 0.23. Table S3), we pooled the abundance data for both species into a single "total stickleback" metric for the analysis of light trap efficiency. In the analysis of stickleback catch in benthic traps, we excluded *P. pungitius* from the analysis due to its low frequency of occurrence (17.7 %, Table S4) which created a zero-inflated distribution that precluded robust parameter estimation for environmental drivers. Consequently, we restricted the final mechanistic assessment of temperature versus night duration in benthic traps to the dominant species, *G. aculeatus*, which was present in 74.7% of hauls (Table S4).

Based on these preliminary tests, we fitted final separate GLMMs for *N. integer*, *P. flexuosus*, pooled sticklebacks (light traps), and *G. aculeatus* (benthic traps). The fixed effects for all light trap models included *Trap type* (Modified net vs. Unmodified) and *Night duration*. For the benthic trap *G. aculeatus* model, *Water temperature* was included as the only environmental covariate (Eq. S2).

To test the hypothesis that stickleback presence negatively affects mysid catches, whether through direct predation or chemically/visually mediated avoidance, we fitted an additional model for *N. integer*. We restricted this biotic interaction analysis to *N. integer* because it was the numerically dominant species (77% of total catch in modified light traps, Table S4). Although both species occurred with similar frequency, *P. flexuosus* appeared in significantly lower densities (averaging ~5 individuals per positive catch compared to ~18 for *N. integer*), limiting the statistical resolution required to detect subtle density-dependent responses to predation risk.

In this analysis, stickleback abundance was defined as the combined total of three-spined (*G. aculeatus*) and ninespine (*P. pungitius*) sticklebacks per trap. We pooled these species under the assumption that both contribute to the local “risk landscape” by emitting olfactory cues (kairomones) that trigger anti-predator behavior in mysids (Hamrén and Hansson 1999; Lindén, Lehtiniemi and Viitasalo 2003). This pooled abundance was treated as a fixed predictor, controlling for *Night duration* (Eq. 3).

To evaluate whether potential predator effects were specific to sticklebacks or a general consequence of high fish densities, we fitted an identical model replacing stickleback abundance with the total abundance of pipefish (*Nerophis ophidion* and *Syngnathus typhle*). Syngnathids are also common in these littoral habitats and are capable of preying on mysids (Van Wassenbergh *et al.* 2007; Sundin *et al.* 2011), making them a relevant control group to assess inter-specific interaction effects.

Due to the geographical closeness of the traps, we tested for spatial autocorrelation in the model residuals using Moran's I (DHARMa package; Hartig 2022). While minor autocorrelation was detected in one bay (*P. flexuosus* model, Area C, p = 0.03, Table S5), it was absent in the other three areas. Attempts to include a formal spatial covariance structure resulted in model convergence failure, indicating the data were insufficient to support the additional complexity. Therefore, we retained the nested random effect structure (*Station* within *Area*) as the most parsimonious and robust approach.

Model assumptions were verified by inspection of scaled residuals for uniformity, outliers, and overdispersion. No significant deviations from model assumptions were detected in the final models. Significance of fixed effects was assessed using Wald z-tests. Estimated marginal means and confidence intervals were extracted using the sjPlot package (Lüdecke 2025) for visualization. Model coefficients were exponentiated to derive Incidence Rate Ratios (IRR) and their associated 95% confidence intervals. An IRR of 1 indicates no change in catch rate, while values < 1 or > 1 indicate a decrease or increase relative to the reference level, respectively.

$$\text{count}_{ij}\sim\text{NB}\left( \mu_{ij},\theta\right)$$

$$\log\left( \mu_{ij} \right)=\beta_{0}+\beta_{1}\text{Species}_{ij}+\beta_{2}\text{Trap type}_{ij}+\beta_{3}\text{Night duration}_{ij}+\beta_{4}\left( \text{Species}\times\text{Trap type} \right)_{ij}+\beta_{5}\left( \text{Species}\times\text{Night duration} \right)_{ij}$$

$$+ u_{j}u_{j}\sim N\left( 0,\sigma_{\text{Station(Area)}}^{2} \right)$$

Eq. S1

$$\begin{matrix} \text{count}_{ij}\sim\text{NB}\left( \mu_{ij},\theta\right) \\ \log\left( \mu_{ij} \right)=\beta_{0}+\beta_{1}\text{Water temperature}_{ij} \end{matrix}$$

$$+ u_{j}u_{j}\sim N\left( 0,\sigma_{\text{Station(Area)}}^{2} \right)$$

Eq. S2

$$\begin{matrix} \text{count}_{ij}\sim\text{NB}\left( \mu_{ij},\theta\right) \\ \log\left( \mu_{ij} \right)=\beta_{0}+\beta_{1}\text{Stickleback abundance}_{ij}+ \beta_{1}\text{Night duration}_{ij} \end{matrix}$$

$$+ u_{j}u_{j}\sim N\left( 0,\sigma_{\text{Station(Area)}}^{2} \right)$$

Eq. S3

#### S5. Precision optimization, power analysis procedure and parameterization

##### *Precision optimization*

Using a Monte Carlo approach, we simulated random draws (n = 10,000) from negative binomial distributions parameterized by the empirical mean density (µ) and dispersion (θ) specific to *N. integer* and pooled sticklebacks (*G. aculeatus* and *P. pungitius*). We calculated the reduction in relative standard error (CV) as sample size increased from 1 to 30 traps. The optimal sample size was defined as the point where the rate of precision improvement (reduction in CV) dropped below 20% of the initial rate.

##### *Power analysis simulations*

We performed a hierarchical power analysis to determine the method's sensitivity to population changes while accounting for spatial variability. We simulated a monitoring program across four areas with a fixed sampling effort of 10 traps per area, matching the optimal effort identified in the precision analysis. Unlike the precision analysis, count data were generated using a mixed-effects simulation structure (Eq. S1) to explicitly model the random variation between areas ($\sigma_{Area}$) alongside the specific overdispersion properties (θ) of each functional group.

For *N. integer*, we utilized parameters derived from modified traps. To strictly isolate the influence of aggregation on detection power, we standardized the baseline mean density for both *N. integer* and the sticklebacks to a fixed low-density spring scenario (μ = 5 ind./trap). This ensured that any differences in power between the highly aggregated mysids (θ ≈ 0.10) and the more uniformly distributed sticklebacks (θ ≈ 0.62) were driven solely by dispersion and spatial variance rather than differences in abundance.

We simulated population changes of varying magnitudes to assess the gear's ability to track both declines and increases in population size. Scenarios included declines ranging from 50% to 90%, as well as population increases (1.5-fold to 5-fold) to simulate recruitment events. To visualize the mechanism limiting statistical power, we also generated sampling distribution plots at the optimal effort (n = 10). These compared the distribution of 1,000 simulated mean catches between the reference and impact (50% decline) scenarios to quantify the degree of overlap masking the population signal.

For each combination of species, sample size, and effect size, we generated 1,000 Monte Carlo datasets. We fitted a negative binomial GLMM to each simulated dataset (Eq. S1), testing for a significant effect of *Year (two-level factor)*. Statistical power was calculated as the proportion of simulations in which the null hypothesis was correctly rejected (Wald z-test, α = 0.05).

$$\begin{matrix} \text{count}_{ij}\sim\text{NB}\left( \mu_{ij},\theta\right) \\ \log\left( \mu_{ij} \right)=\beta_{0}+\beta_{1}\text{Year}_{ij} \end{matrix}$$

$$+ u_{j}u_{j}\sim N\left( 0,\sigma_{Area}^{2} \right)$$

Eq. S1

**Supporting Information Figures**

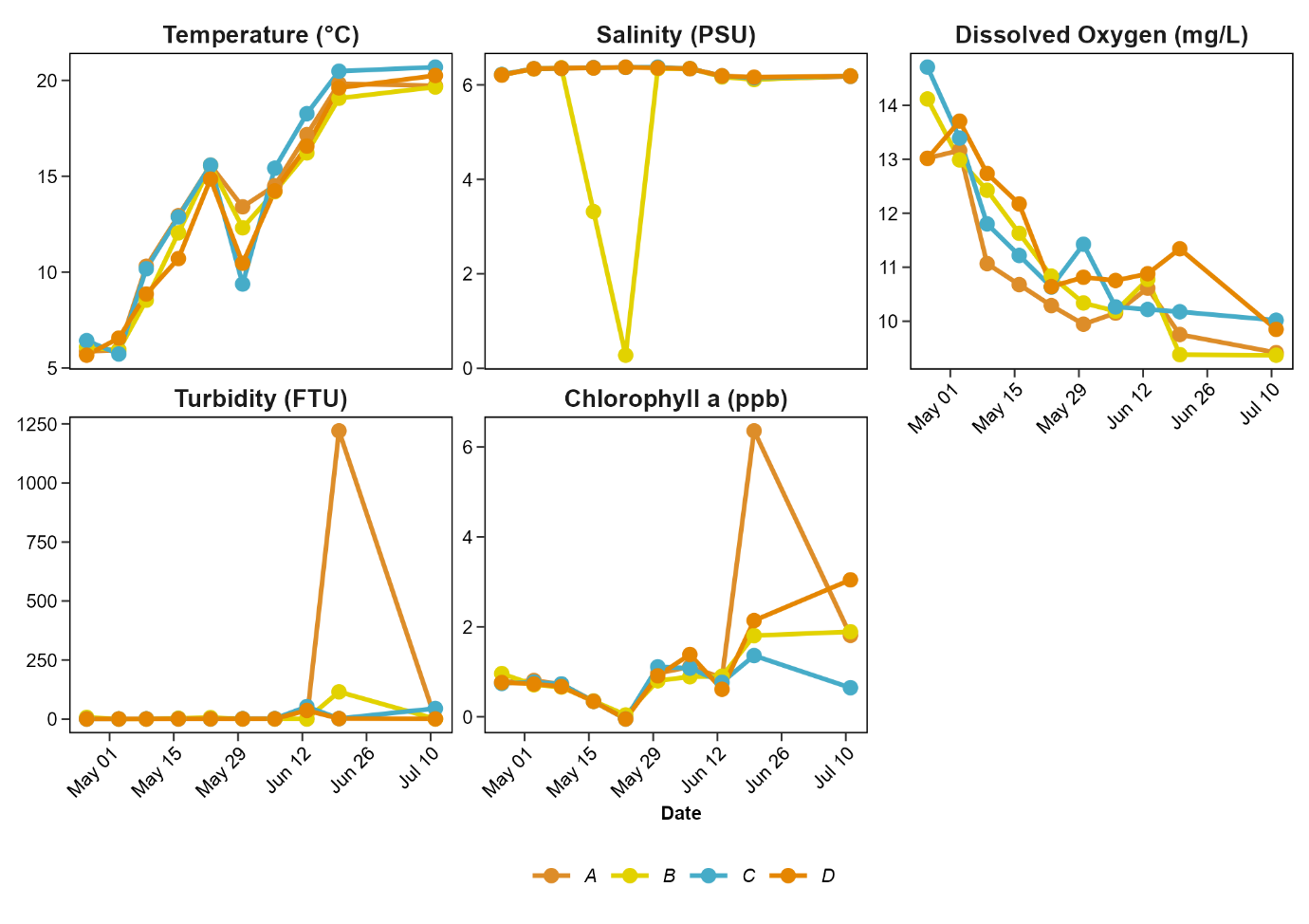

Figure S1

. Temporal variation in environmental variables across the four sampling areas during the study period (April–July). Panels display mean values per sampling date for water temperature (°C), salinity (PSU), dissolved oxygen (mg/L), turbidity (FTU), and chlorophyll a fluorescence (ppb). Coloured lines and points distinguish specific sampling bays (Area).

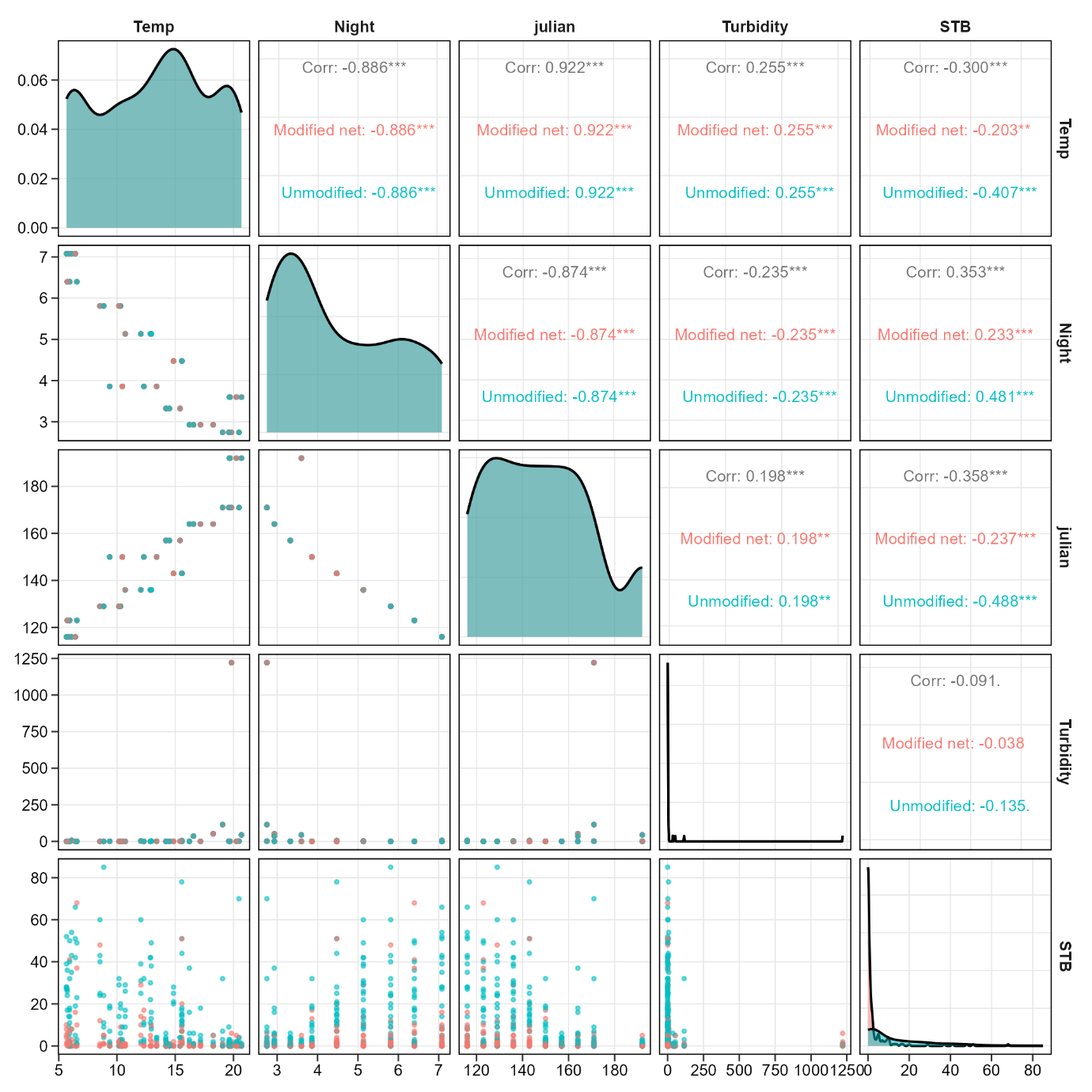

Figure S2

. Correlation matrix of presumptive input variables to mysid models. Temp = Temperature (°C), Night = Night duration h), Julian = Julian date, Turbidity = Turbidity (FTU) and STB = Stickleback abundance (G. aculeatus + P. pungitius)..

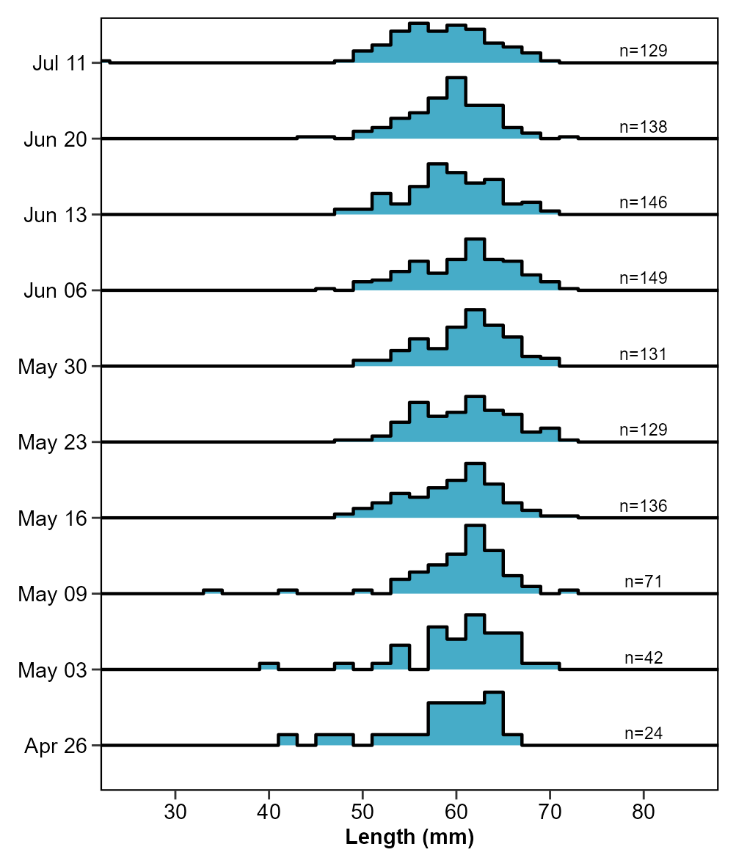

Figure S3

. Seasonal length-frequency distributions of three-spined stickleback (Gasterosteus aculeatus) caught in unlit benthic traps (non-size-selective control). Histograms (bin width = 2 mm) show the relative size structure of the demersal population from April to July. Annotations (n) indicate the total abundance per sampling date.

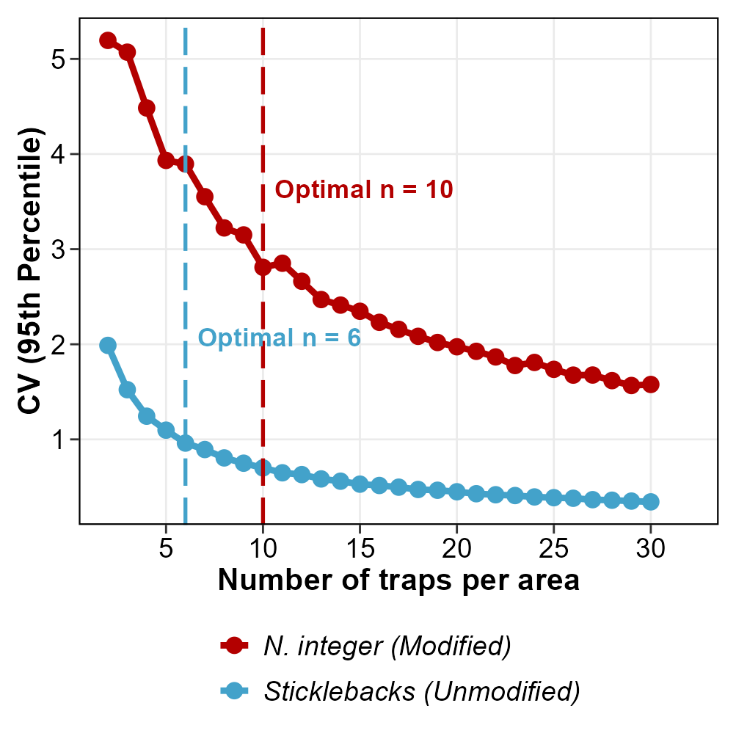

Figure S4

. Relationship between sampling effort and precision for littoral mysids (N. integer) and sticklebacks (pooled G. aculeatus and P. pungitius) showing the 95^th^ percentile of the coefficient of variation distribution as a function of sampling effort (number of traps per area). The dashed vertical lines indicate the optimal sampling effort for each respective group, defined as the point at which the rate of precision improvement dropped below 20% of the initial rate.

**Supporting Information Tables**

Table S1

. Model selection results comparing the explanatory power of Night duration versus Water temperature for determining catch rates. Models were ranked using Akaike Information Criterion (AIC). The predictor yielding the lowest AIC was selected for the final analysis.

| **Species/Group** | **Trap type** | **Model: Night duration (AIC)** | **Model: Temperature (AIC)** | **ΔAIC** | **Selected Driver** |
| --- | --- | --- | --- | --- | --- |
| N. integer | Light trap | 771.7 | 778.5 | 6.8 | Night Duration |
| P. flexuosus | Light trap | 608.1 | 624.1 | 16.1 | Night Duration |
| G. aculeatus + P. pungitius | Light trap | 2103.0 | 2128.4 | 25.4 | Night Duration |

Table S2

. Generalized mixed model results for a combined model including Species (N. integer and P. flexuosus) as a fixed factor and it’s interaction with Trap type and Night duration.

|  | *Estimated Abundance (Count)* | | | |
| --- | --- | --- | --- | --- |
| **Predictors** | **IRR^*^** | **SE** | **Statistic** | **p** |
| Intercept | 1.10 | 0.55 | 0.20 | 0.844 |
| Species (*Praunus flexuosus*) | 0.37 | 0.16 | -2.28 | **0.023** |
| Trap Type (Unmodified) | 0.14 | 0.07 | -3.93 | **<0.001** |
| Night duration (centered) | 1.48 | 0.22 | 2.64 | **0.008** |
| Species × Trap type | 0.71 | 0.48 | -0.51 | 0.612 |
| Species × Night duration | 2.11 | 0.58 | 2.72 | **0.006** |
| **Random Effects** | | | | |
| σ^2^ | 2.33 | | | |
| τ_00_ _Station:Area_ | 4.01 | | | |
| τ_00_ _Area_ | 0.19 | | | |
| ICC | 0.64 | | | |
| N _Station_ | 10 | | | |
| N _Area_ | 4 | | | |
| Observations | 800 | | | |
| Marginal R^2^ / Conditional R^2^ | 0.311 / 0.754 | | | |
| * IRR = Incidence Rate Ratio |  | | | |

Table S3

. Generalized linear mixed model results for species-specific responses (sticklebacks) to environmental variables in light traps.

| **Predictors** | **IRR** | **SE** | **Statistic** | **p** |
| --- | --- | --- | --- | --- |
| Intercept | 2.20 | 0.45 | 3.82 | **<0.001** |
| Night duration (centered) | 2.49 | 0.43 | 5.24 | **<0.001** |
| Species: TSS (vs NSS) | 1.68 | 0.26 | 3.34 | **0.001** |
| Temperature (centered) | 1.07 | 0.06 | 1.20 | 0.228 |
| Night duration × Species (TSS) | 0.74 | 0.19 | -1.20 | 0.229 |
| Temp × Species (TSS) | 0.97 | 0.07 | -0.40 | 0.693 |
| **Random Effects** | | | | |
| σ^2^ | 1.66 | | | |
| τ_00_ _Area:Station_ | 0.22 | | | |
| ICC | 0.12 | | | |
| N _Area_ | 4 | | | |
| N _Station_ | 10 | | | |
| Observations | 800 | | | |
| Marginal R^2^ / Conditional R^2^ | 0.317 / 0.398 | | | |
| TSS = Three-spined stickleback (*G. aculateus*),  NSS = Ninespine stickleback (*P. pungitius*)  IRR = Incidence Rate Ratio |  | | | |

Table S4

. Catch summary and prevalence of mysids and sticklebacks across sampling gears. “Total traps” denotes the sampling effort (n). Prevalence indicates the number of samples containing at least one individual of the species and the corresponding frequency of occurrence (%). Percent catch reflects the relative abundance of each species within the total catch for that gear type.

| **Trap type** | **Species** | **Total traps (n)** | **Prevalence (n traps)** | **Freq. occurrence (%)** | **Individuals (n)** | **Percent catch (%)** |
| --- | --- | --- | --- | --- | --- | --- |
| Light trap (Modified) | *N. integer* | 200 | 44 | 22 | 795 | 77.3 |
| Light trap (Modified) | *P. flexuosus* | 200 | 50 | 25 | 234 | 22.7 |
| Benthic trap | *G. aculeatus* | 395 | 70 | 82.3 | 2523 | 93.9 |
| Benthic trap | *P. pungitius* | 395 | 295 | 17.7 | 165 | 6.14 |

Table S5

. Spatial autocorrelation in GLMM residuals.

| **Species** | **Area** | **Moran’s I** | **p-value** |
| --- | --- | --- | --- |
| *N. integer* | A | -0.05 | 0.46 |
| *N. integer* | B | -0.17 | 0.55 |
| *N. integer* | C | 0.09 | 0.03 |
| *N. integer* | D | -0.25 | 0.28 |
| *P. flexuosus* | A | 0.15 | **0.01** |
| *P. flexuosus* | B | -0.16 | 0.65 |
| *P. flexuosus* | C | -0.02 | 0.35 |
| *P. flexuosus* | D | -0.14 | 0.77 |

Table S6

. Catch composition (total abundance) in light traps presented by sampling date, fishing area and trap type. Species are ordered by highest total abundance from left to right. Ner. Oph = Nerophis Ophidion, Gas. acu. = Gasterosteus aculeatus, Pun. pun. = Pungitius pungitius, Neo. Int. = Neomysis integer, Pra. Fle. = Praunus flexuosus, Pal./Cran. = Palaemon adspersus/ Crangon crangon, Syn. typ. = Syngnathus typhle, Larvae = Unidentified fish larvae, Pom. min. = Pomatoschistus minutus, Zoa. viv = Zoarces viviparus, Gob. fla. = Gobiusculus flavescens, Pho. gun. = Pholis gunnellus, Per. flu = Perca fluviatilis.

| **Date** | **Area** | **Trap type** | **Ner. oph.** | **Gas. acu.** | **Pun. pun** | **Neo. int.** | **Pra. fle..** | **Pal./Cran.** | **Syn. typ.** | **Larvae** | **Pom. min.** | **Zoa. viv.** | **Gob. fla.** | **Pho. gun.** | **Per. flu.** |
| --- | --- | --- | --- | --- | --- | --- | --- | --- | --- | --- | --- | --- | --- | --- | --- |
| 2023-04-26 | A | Modified net | 0 | 0 | 1 | 57 | 5 | 0 | 0 | 0 | 0 | 0 | 0 | 0 | 0 |
| 2023-04-26 | A | Unmodified | 0 | 15 | 15 | 99 | 2 | 0 | 0 | 0 | 0 | 0 | 0 | 0 | 0 |
| 2023-04-26 | B | Modified net | 0 | 4 | 7 | 1 | 5 | 0 | 0 | 0 | 0 | 0 | 0 | 0 | 0 |
| 2023-04-26 | B | Unmodified | 0 | 66 | 74 | 3 | 3 | 0 | 0 | 0 | 1 | 0 | 1 | 0 | 0 |
| 2023-04-26 | C | Modified net | 0 | 15 | 3 | 36 | 3 | 0 | 0 | 0 | 0 | 0 | 0 | 0 | 0 |
| 2023-04-26 | C | Unmodified | 0 | 156 | 27 | 0 | 0 | 0 | 0 | 0 | 0 | 0 | 0 | 0 | 0 |
| 2023-04-26 | D | Modified net | 0 | 13 | 15 | 5 | 8 | 0 | 0 | 0 | 0 | 0 | 0 | 0 | 0 |
| 2023-04-26 | D | Unmodified | 0 | 97 | 77 | 1 | 0 | 0 | 0 | 0 | 0 | 0 | 0 | 0 | 0 |
| 2023-05-03 | A | Modified net | 9 | 2 | 0 | 0 | 0 | 0 | 0 | 0 | 0 | 0 | 0 | 0 | 0 |
| 2023-05-03 | A | Unmodified | 3 | 103 | 10 | 0 | 0 | 0 | 0 | 0 | 0 | 0 | 0 | 0 | 0 |
| 2023-05-03 | B | Modified net | 52 | 23 | 30 | 0 | 0 | 0 | 7 | 0 | 0 | 0 | 0 | 0 | 0 |
| 2023-05-03 | B | Unmodified | 35 | 75 | 2 | 0 | 0 | 0 | 12 | 0 | 0 | 0 | 0 | 0 | 0 |
| 2023-05-03 | C | Modified net | 6 | 9 | 1 | 0 | 5 | 0 | 0 | 0 | 0 | 0 | 0 | 0 | 0 |
| 2023-05-03 | C | Unmodified | 3 | 110 | 20 | 0 | 0 | 0 | 0 | 0 | 0 | 0 | 0 | 0 | 0 |
| 2023-05-03 | D | Modified net | 11 | 78 | 37 | 0 | 1 | 0 | 5 | 0 | 0 | 0 | 0 | 0 | 0 |
| 2023-05-03 | D | Unmodified | 14 | 75 | 2 | 0 | 1 | 0 | 1 | 0 | 0 | 0 | 0 | 0 | 0 |
| 2023-05-09 | A | Modified net | 20 | 3 | 9 | 151 | 2 | 0 | 0 | 0 | 0 | 0 | 0 | 0 | 0 |
| 2023-05-09 | A | Unmodified | 34 | 23 | 27 | 1 | 1 | 0 | 0 | 0 | 0 | 0 | 0 | 0 | 0 |
| 2023-05-09 | B | Modified net | 78 | 43 | 18 | 56 | 29 | 0 | 8 | 0 | 0 | 0 | 0 | 0 | 0 |
| 2023-05-09 | B | Unmodified | 125 | 83 | 109 | 0 | 0 | 0 | 11 | 0 | 0 | 0 | 0 | 0 | 0 |
| 2023-05-09 | C | Modified net | 44 | 4 | 3 | 10 | 27 | 0 | 1 | 0 | 0 | 0 | 0 | 0 | 0 |
| 2023-05-09 | C | Unmodified | 31 | 73 | 2 | 0 | 10 | 0 | 7 | 0 | 0 | 0 | 0 | 0 | 0 |
| 2023-05-09 | D | Modified net | 50 | 8 | 5 | 5 | 13 | 0 | 4 | 3 | 0 | 0 | 0 | 0 | 0 |
| 2023-05-09 | D | Unmodified | 58 | 81 | 52 | 2 | 11 | 0 | 10 | 0 | 0 | 0 | 0 | 0 | 0 |
| 2023-05-16 | A | Modified net | 8 | 0 | 21 | 34 | 16 | 0 | 1 | 1 | 0 | 0 | 0 | 0 | 0 |
| 2023-05-16 | A | Unmodified | 10 | 34 | 90 | 0 | 0 | 0 | 0 | 0 | 0 | 0 | 0 | 0 | 0 |
| 2023-05-16 | B | Modified net | 31 | 14 | 45 | 17 | 50 | 0 | 1 | 3 | 2 | 1 | 0 | 0 | 0 |
| 2023-05-16 | B | Unmodified | 85 | 95 | 62 | 0 | 0 | 0 | 0 | 0 | 0 | 0 | 0 | 0 | 0 |
| 2023-05-16 | C | Modified net | 107 | 1 | 7 | 135 | 20 | 0 | 5 | 3 | 0 | 1 | 0 | 0 | 0 |
| 2023-05-16 | C | Unmodified | 94 | 109 | 38 | 59 | 13 | 0 | 11 | 0 | 1 | 0 | 0 | 0 | 0 |
| 2023-05-16 | D | Modified net | 96 | 1 | 1 | 3 | 21 | 0 | 12 | 0 | 2 | 1 | 0 | 0 | 0 |
| 2023-05-16 | D | Unmodified | 220 | 59 | 36 | 0 | 0 | 0 | 10 | 0 | 0 | 2 | 0 | 0 | 0 |
| 2023-05-23 | A | Modified net | 15 | 16 | 26 | 16 | 5 | 0 | 0 | 3 | 1 | 0 | 0 | 0 | 0 |
| 2023-05-23 | A | Unmodified | 2 | 19 | 62 | 1 | 1 | 0 | 0 | 0 | 1 | 0 | 0 | 0 | 0 |
| 2023-05-23 | B | Modified net | 41 | 2 | 20 | 0 | 6 | 0 | 0 | 0 | 0 | 1 | 0 | 0 | 0 |
| 2023-05-23 | B | Unmodified | 55 | 84 | 70 | 12 | 45 | 0 | 0 | 0 | 0 | 0 | 0 | 0 | 0 |
| 2023-05-23 | C | Modified net | 0 | 2 | 54 | 131 | 12 | 0 | 0 | 0 | 0 | 0 | 0 | 0 | 0 |
| 2023-05-23 | C | Unmodified | 0 | 12 | 60 | 69 | 2 | 0 | 0 | 0 | 0 | 0 | 0 | 0 | 0 |
| 2023-05-23 | D | Modified net | 131 | 0 | 2 | 0 | 1 | 0 | 1 | 1 | 0 | 1 | 0 | 0 | 0 |
| 2023-05-23 | D | Unmodified | 50 | 77 | 17 | 0 | 0 | 0 | 4 | 0 | 0 | 1 | 0 | 0 | 0 |
| 2023-05-30 | A | Modified net | 60 | 1 | 2 | 0 | 0 | 0 | 0 | 0 | 0 | 0 | 0 | 0 | 0 |
| 2023-05-30 | A | Unmodified | 21 | 9 | 2 | 0 | 0 | 1 | 0 | 0 | 0 | 0 | 0 | 0 | 0 |
| 2023-05-30 | B | Modified net | 0 | 1 | 30 | 6 | 3 | 0 | 0 | 0 | 0 | 0 | 0 | 0 | 0 |
| 2023-05-30 | B | Unmodified | 0 | 4 | 54 | 0 | 12 | 0 | 0 | 1 | 2 | 0 | 0 | 0 | 0 |
| 2023-05-30 | C | Modified net | 48 | 1 | 1 | 0 | 0 | 0 | 0 | 0 | 0 | 0 | 0 | 0 | 0 |
| 2023-05-30 | C | Unmodified | 72 | 29 | 17 | 0 | 0 | 0 | 2 | 0 | 0 | 0 | 0 | 0 | 0 |
| 2023-05-30 | D | Modified net | 124 | 0 | 1 | 0 | 0 | 0 | 1 | 0 | 0 | 0 | 0 | 0 | 0 |
| 2023-05-30 | D | Unmodified | 203 | 10 | 4 | 0 | 0 | 0 | 0 | 0 | 0 | 0 | 0 | 0 | 0 |
| 2023-06-06 | A | Modified net | 7 | 1 | 0 | 0 | 0 | 0 | 0 | 0 | 0 | 0 | 0 | 0 | 0 |
| 2023-06-06 | A | Unmodified | 5 | 1 | 0 | 0 | 0 | 0 | 0 | 0 | 0 | 0 | 0 | 0 | 0 |
| 2023-06-06 | B | Modified net | 4 | 0 | 2 | 0 | 0 | 0 | 0 | 0 | 0 | 0 | 0 | 0 | 0 |
| 2023-06-06 | B | Unmodified | 2 | 6 | 10 | 0 | 1 | 0 | 0 | 0 | 0 | 0 | 0 | 0 | 0 |
| 2023-06-06 | C | Modified net | 11 | 2 | 0 | 0 | 1 | 0 | 0 | 0 | 0 | 0 | 0 | 0 | 0 |
| 2023-06-06 | C | Unmodified | 12 | 5 | 3 | 0 | 0 | 0 | 1 | 0 | 0 | 0 | 0 | 0 | 0 |
| 2023-06-06 | D | Modified net | 40 | 1 | 0 | 0 | 0 | 0 | 0 | 0 | 0 | 0 | 0 | 1 | 0 |
| 2023-06-06 | D | Unmodified | 56 | 3 | 0 | 0 | 0 | 0 | 2 | 0 | 0 | 0 | 0 | 0 | 0 |
| 2023-06-13 | A | Modified net | 2 | 0 | 1 | 11 | 1 | 0 | 0 | 0 | 0 | 0 | 0 | 0 | 0 |
| 2023-06-13 | A | Unmodified | 2 | 28 | 1 | 4 | 0 | 0 | 0 | 0 | 0 | 0 | 0 | 0 | 0 |
| 2023-06-13 | B | Modified net | 0 | 0 | 1 | 0 | 0 | 0 | 0 | 13 | 0 | 0 | 0 | 0 | 0 |
| 2023-06-13 | B | Unmodified | 0 | 56 | 8 | 0 | 0 | 0 | 0 | 0 | 0 | 0 | 1 | 0 | 0 |
| 2023-06-13 | C | Modified net | 11 | 3 | 2 | 0 | 0 | 0 | 0 | 0 | 0 | 0 | 0 | 0 | 0 |
| 2023-06-13 | C | Unmodified | 5 | 7 | 1 | 0 | 0 | 24 | 2 | 1 | 0 | 0 | 0 | 0 | 0 |
| 2023-06-13 | D | Modified net | 9 | 1 | 1 | 1 | 0 | 5 | 0 | 2 | 0 | 0 | 0 | 0 | 0 |
| 2023-06-13 | D | Unmodified | 8 | 7 | 1 | 0 | 0 | 36 | 1 | 0 | 0 | 0 | 0 | 0 | 0 |
| 2023-06-20 | A | Modified net | 1 | 8 | 0 | 64 | 0 | 0 | 0 | 0 | 0 | 0 | 0 | 0 | 0 |
| 2023-06-20 | A | Unmodified | 0 | 2 | 1 | 32 | 0 | 6 | 0 | 1 | 0 | 0 | 0 | 0 | 1 |
| 2023-06-20 | B | Modified net | 0 | 0 | 1 | 0 | 0 | 2 | 0 | 1 | 0 | 0 | 0 | 0 | 0 |
| 2023-06-20 | B | Unmodified | 0 | 40 | 3 | 0 | 0 | 8 | 0 | 1 | 0 | 0 | 0 | 0 | 0 |
| 2023-06-20 | C | Modified net | 1 | 5 | 0 | 0 | 0 | 12 | 0 | 0 | 0 | 0 | 0 | 0 | 0 |
| 2023-06-20 | C | Unmodified | 1 | 77 | 0 | 0 | 0 | 35 | 0 | 0 | 0 | 0 | 0 | 0 | 0 |
| 2023-06-20 | D | Modified net | 2 | 2 | 1 | 0 | 0 | 9 | 0 | 1 | 0 | 0 | 0 | 0 | 0 |
| 2023-06-20 | D | Unmodified | 1 | 10 | 2 | 0 | 0 | 52 | 1 | 0 | 0 | 0 | 0 | 1 | 0 |
| 2023-07-11 | A | Modified net | 0 | 0 | 0 | 56 | 0 | 0 | 0 | 1 | 0 | 0 | 0 | 0 | 0 |
| 2023-07-11 | A | Unmodified | 1 | 3 | 0 | 0 | 0 | 0 | 0 | 0 | 0 | 0 | 0 | 0 | 0 |
| 2023-07-11 | B | Modified net | 1 | 0 | 0 | 0 | 0 | 0 | 0 | 15 | 0 | 0 | 0 | 0 | 0 |
| 2023-07-11 | B | Unmodified | 1 | 7 | 0 | 0 | 0 | 0 | 1 | 0 | 0 | 0 | 0 | 0 | 0 |
| 2023-07-11 | C | Modified net | 0 | 0 | 0 | 0 | 0 | 0 | 0 | 43 | 0 | 0 | 0 | 0 | 0 |
| 2023-07-11 | C | Unmodified | 1 | 2 | 0 | 0 | 0 | 0 | 0 | 19 | 0 | 0 | 0 | 0 | 0 |
| 2023-07-11 | D | Modified net | 2 | 5 | 0 | 0 | 0 | 2 | 0 | 7 | 0 | 0 | 0 | 0 | 0 |
| 2023-07-11 | D | Unmodified | 2 | 5 | 0 | 0 | 0 | 0 | 0 | 0 | 0 | 0 | 0 | 0 | 0 |

Table S7

. Model selection for environmental drivers of mysid abundance in light traps. Comparison of Night duration vs. Temperature models.

|  | **Neomysis (Night)** | | | | **Neomysis (Temp)** | | | | **Praunus (Night)** | | | | **Praunus (Temp)** | | | |
| --- | --- | --- | --- | --- | --- | --- | --- | --- | --- | --- | --- | --- | --- | --- | --- | --- |
| *Predictors* | *Log-Mean* | *SE* | *Statistic* | *p* | *Log-Mean* | *SE* | *Statistic* | *p* | *Log-Mean* | *SE* | *Statistic* | *p* | *Log-Mean* | *SEr* | *Statistic* | *p* |
| Intercept | -0.07 | 0.91 | -0.08 | 0.934 | 0.16 | 0.89 | 0.18 | 0.857 | -1.09 | 0.41 | -2.67 | **0.008** | -0.81 | 0.40 | -2.01 | **0.045** |
| Trap type (Unmodified) | -1.92 | 0.60 | -3.20 | **0.001** | -1.95 | 0.60 | -3.22 | **0.001** | -1.73 | 0.46 | -3.74 | **<0.001** | -1.76 | 0.47 | -3.74 | **<0.001** |
| Night duration (centered) | 0.54 | 0.19 | 2.89 | **0.004** |  |  |  |  | 0.95 | 0.18 | 5.13 | **<0.001** |  |  |  |  |
| Temperature (centered) |  |  |  |  | -0.08 | 0.05 | -1.56 | 0.118 |  |  |  |  | -0.20 | 0.05 | -3.69 | **<0.001** |
| **Random Effects** | | | | | | | | | | | | | | | | |
| σ^2^ | 2.49 | | | | 2.57 | | | | 2.01 | | | |  |  |  |  |
| τ_00_ | 2.80 _Station:Area_ | | | | 2.48 _Station:Area_ | | | | 2.71 _Station:Area_ | | | | 2.65 _Station:Area_ | | | |
|  | 2.39 _Area_ | | | | 2.32 _Area_ | | | | 0.00 _Area_ | | | | 0.00 _Area_ | | | |
| ICC | 0.68 | | | | 0.65 | | | | 0.57 | | | |  | | | |
| N | 10 _Station_ | | | | 10 _Station_ | | | | 10 _Station_ | | | | 10 _Station_ | | | |
|  | 4 _Area_ | | | | 4 _Area_ | | | | 4 _Area_ | | | | 4 _Area_ | | | |
| Observations | 400 | | | | 400 | | | | 400 | | | | 400 | | | |
| Marginal R^2^ / Conditional R^2^ | 0.165 / 0.729 | | | | 0.132 / 0.697 | | | | 0.355 / 0.725 | | | | NA | | | |
| AIC | 771.657 | | | | 778.495 | | | | 608.065 | | | | 624.126 | | | |

Table S8

. Model selection for benthic trap catches of G. aculeatus. The continuous predictors Temperature and Night duration have been mean-centered.

|  | **Full Model** | | | | | | | **Night Only** | | | | | | | | **Temp Only** | | | | | | | |
| --- | --- | --- | --- | --- | --- | --- | --- | --- | --- | --- | --- | --- | --- | --- | --- | --- | --- | --- | --- | --- | --- | --- | --- |
| *Predictors* | *IRR* | *SE* | | *Statistic* | | *p* | | *IRR* | | *SE* | | *Statistic* | | *p* | | *IRR* | | *SE* | | *Statistic* | | *p* | |
| Intercept | 4.31 | | 0.57 | | 11.07 | | **<0.001** | | 4.33 | | 0.56 | | 11.36 | | **<0.001** | | 4.47 | | 0.65 | | 10.24 | | **<0.001** |
| Temperature (centered) | 1.04 | | 0.03 | | 1.12 | | 0.261 | |  | |  | |  | |  | | 1.18 | | 0.02 | | 9.45 | | **<0.001** |
| Night duration (centered) | 0.63 | | 0.07 | | -4.31 | | **<0.001** | | 0.56 | | 0.03 | | -10.70 | | **<0.001** | |  | |  | |  | |  |
| **Random Effects** | | | | | | | | | | | | | | | | | | | | | | | |
| σ^2^ | 0.97 | | | | | | | 0.97 | | | | | | | | 1.00 | | | | | | | |
| τ_00_ | 0.20 _Station:Area_ | | | | | | | 0.19 _Station:Area_ | | | | | | | | 0.24 _Station:Area_ | | | | | | | |
|  | 0.03 _Area_ | | | | | | | 0.03 _Area_ | | | | | | | | 0.04 _Area_ | | | | | | | |
| ICC | 0.19 | | | | | | | 0.18 | | | | | | | | 0.22 | | | | | | | |
| N | 10 _Station_ | | | | | | | 10 _Station_ | | | | | | | | 10 _Station_ | | | | | | | |
|  | 4 _Area_ | | | | | | | 4 _Area_ | | | | | | | | 4 _Area_ | | | | | | | |
| Observations | 395 | | | | | | | 395 | | | | | | | | 395 | | | | | | | |
| Marginal R^2^ / Conditional R^2^ | 0.365 / 0.487 | | | | | | | 0.361 / 0.478 | | | | | | | | 0.326 / 0.475 | | | | | | | |
| AIC | 2082.360 | | | | | | | 2081.621 | | | | | | | | 2098.975 | | | | | | | |
| IRR = Incidence Rate Ratio |  | | | | | | |  | | | | | | | |  | | | | | | | |

### Supporting Information References

Adams CM, Jones MT, Hatch R *et al.* Laboratory and field evaluation of a new light‐emitting‐diode‐lit quatrefoil light trap for sampling fish early life stages. *North American Journal of Fisheries Management* 2023;**43**:1337–48.

Boscarino BT, Rudstam LG, Loew ER *et al.* Predicting the vertical distribution of the opossum shrimp, Mysis relicta, in Lake Ontario: a test of laboratory-based light preferences. *Canadian Journal of Fisheries and Aquatic Sciences* 2009;**66**:101–13.

Hamrén U, Hansson S. A mysid shrimp (Mysis mixta) is able to detect the odour of its predator (Clupea harengus). *Ophelia* 1999;**51**:187–91.

Kyba CCM, Mohar A, Posch T. How bright is moonlight? *A&G* 2017;**58**:1.31-1.32.

Lindén E, Lehtiniemi M, Viitasalo M. Predator avoidance behaviour of Baltic littoral mysids Neomysis integer and Praunus flexuosus. *Marine Biology* 2003;**143**:845–50.

Lindström M. Spectral sensitivity and light tolerance of mysid species in the Baltic area. In: Köhn J, Jones MB, Moffat A (eds), *Taxonomy, Biology and Ecology of (Baltic) Mysids (Mysidacea: Crustacea)*. 1st edn. Hiddensee, Germany: Rostock University, 1992, 120–6.

Lüdecke D. *sjPlot: Data Visualization for Statistics in Social Science*., 2025.

Sundin J, Jacobsson Ö, Berglund A *et al.* Straight-nosed pipefish Nerophis ophidion and broad-nosed pipefish Syngnathus typhle avoid eelgrass overgrown with filamentous algae. *Journal of Fish Biology* 2011;**78**:1855–60.

Van Wassenbergh S, Strother JA, Flammang BE *et al.* Extremely fast prey capture in pipefish is powered by elastic recoil. *J R Soc Interface* 2007;**5**:285–96.
