## Supplementary material for "Optimizing light traps for littoral mysids and mesopredatory fish in the Baltic Sea: environmental drivers and seasonal monitoring efficacy": Data and R-code: Analysis_script_PeerJ.R

```
# ==============================================================================
# SCRIPT: Light Trap & Fish Trap Analysis (Full Manuscript & SI)
# JOURNAL: PeerJ
# MANUSCRIPT TITLE: "Optimizing Light Traps for Littoral Mysids and Mesopredatory Fish in the Baltic Sea: Environmental Drivers and Seasonal Monitoring Efficacy"
# AUTHORS: Martin Ogonowski
# INSTITUTE: Swedish University of Agricultural Sciences
# DEPARTMENT: Department of Aquatic Resources, Institute of Freshwater Research, Sweden
# EMAIL:
# DATE: 2026-02-12
# DESCRIPTION: 
#    Reproduces ALL Tables (1-2, S1-S8) and Figures (4-9, S1-S4).
#    
# ==============================================================================

# --- 1. SETUP & LIBRARIES ---
library(tidyverse)
library(data.table)
library(lubridate)
library(glmmTMB)      
library(suncalc)      
library(DHARMa)       
library(sjPlot)       
library(wesanderson)  
library(patchwork)    
library(ggridges)     
library(flextable)    
library(GGally)       
library(scales)
library(officer)

set.seed(42)
Sys.setlocale("LC_TIME", "C") 

# --- CUSTOM THEME (7pt) ---
theme_custom <- function(base_size = 7, base_family = "sans") {
  theme_bw(base_size = base_size, base_family = base_family) +
    theme(
      axis.text = element_text(size = 7, color = "black"),
      axis.title = element_text(size = 7, face = "bold"),
      axis.ticks.length = unit(0.1, "cm"),
      panel.grid = element_blank(),
      panel.border = element_rect(colour = "black", fill = NA, linewidth = 0.5),
      legend.position = "bottom",
      legend.text = element_text(size = 7, face = "italic"),
      legend.title = element_blank(),
      strip.background = element_blank(),
      strip.text = element_text(face = "bold", size = 7),
      plot.tag = element_text(face = "bold", size = 8),
      plot.margin = margin(2, 2, 2, 2, "mm")
    )
}

cols <- wes_palette("FantasticFox1")
col_neo <- cols[5] 
col_pra <- cols[1] 
col_stb <- cols[3] 

# ==============================================================================
# 2. DATA LOADING & PREP
# ==============================================================================

# Load Cleaned Data (Semicolon separated, UTF-8)
lighttrap  <- fread("Clean_data/Lighttrap_clean.csv", sep=";", stringsAsFactors=T, encoding="UTF-8")
stations   <- fread("Clean_data/Stations_clean.csv", sep=";", encoding="UTF-8")
mysid_size <- fread("Clean_data/Mysid_size_clean.csv", sep=";", encoding="UTF-8")
fishtrap   <- fread("Clean_data/Fishtrap_clean.csv", sep=";", encoding="UTF-8")
stb_size   <- fread("Clean_data/Stickleback_size_clean.csv", sep=";", encoding="UTF-8")

# Prep Light Trap Data
lighttrap$Station_comb <- paste0(lighttrap$Bay_ID, lighttrap$Station)
lighttrap$TRIP <- as.factor(lighttrap$TRIP)

# Calculate Night Duration
dates <- data.frame(date=as.Date(unique(lighttrap$Date_Out)), lat=58.82, lon=17.63)
sun <- getSunlightTimes(data=dates, tz="CET") %>% 
  mutate(night2 = as.numeric(24 - (dusk - dawn)), Date_Out = as.character(date)) %>% 
  select(Date_Out, night2)

# Merge
lighttrap$Date_Out <- as.character(lighttrap$Date_Out)
df <- left_join(lighttrap, sun, by="Date_Out")
df <- left_join(df, stations[,.(Station_ID, Lat_DD, Long_DD)], by=c("Station_comb"="Station_ID"))

# Centering night and temp
df <- df %>% 
  mutate(
    night_c = as.numeric(scale(night2, scale=FALSE)), 
    temp_c  = as.numeric(scale(Temp_C, scale=FALSE)))

# Add Pooled Stickleback Counts (Covariate for Mysid Models)
stb_cov <- df %>% 
  filter(species %in% c("Gasterosteus aculeatus", "Pungitius pungitius")) %>% 
  group_by(Trap_ID) %>% summarise(count_stb = sum(count, na.rm=T))

df <- left_join(df, stb_cov, by="Trap_ID")
df$count_stb[is.na(df$count_stb)] <- 0

# --- DATA SUBSETS ---
neo <- df %>% 
  filter(species == "Neomysis integer", Trap_Type %in% c("Modified net", "Unmodified")) %>% 
  mutate(
    night_c = as.vector(scale(night2, scale=FALSE)), 
    temp_c  = as.vector(scale(Temp_C, scale=FALSE)), 
    count_stb_c = as.vector(scale(count_stb, scale=FALSE))
  )

pra <- df %>% 
  filter(species == "Praunus flexuosus", Trap_Type %in% c("Modified net", "Unmodified")) %>% 
  mutate(
    night_c = as.vector(scale(night2, scale=FALSE)), 
    temp_c  = as.vector(scale(Temp_C, scale=FALSE)), 
    count_stb_c = as.vector(scale(count_stb, scale=FALSE))
  )

stb_pooled <- df %>% 
  filter(species %in% c("Gasterosteus aculeatus", "Pungitius pungitius"), 
         Trap_Type %in% c("Modified net", "Unmodified")) %>% 
  group_by(TRIP, Date_Out, Bay_ID, Station, Trap_Type, night2) %>% 
  dplyr::summarise(
    count = sum(count), 
    Temp_C = mean(Temp_C, na.rm=T), 
    .groups="drop"
  ) %>% 
  mutate(
    night_c = as.vector(scale(night2, scale=FALSE)),
    temp_c  = as.vector(scale(Temp_C, scale=FALSE))
  )

# Benthic Trap Data
fishtrap$Date_Out <- as.character(fishtrap$Date_Out)
ft_df <- left_join(fishtrap, sun, by="Date_Out")

ft_sts <- ft_df %>% 
  filter(species == "Gasterosteus aculeatus", TRIP != 1) %>% 
  mutate(
    night_c = as.vector(scale(night2, scale=FALSE)), 
    temp_c = as.vector(scale(Temp_C, scale=FALSE))
  )

# Add Pooled Pipefish Counts (Control Covariate for Mysid Models)
pipe_cov <- df %>% 
  filter(species %in% c("Nerophis ophidion", "Syngnathus typhle")) %>% 
  group_by(Trap_ID) %>% 
  summarise(count_pipe = sum(count, na.rm=T))

df <- left_join(df, pipe_cov, by="Trap_ID")
df$count_pipe[is.na(df$count_pipe)] <- 0

# Update the object "neo" to include the centered pipefish count:
neo <- df %>% 
  filter(species == "Neomysis integer", Trap_Type %in% c("Modified net", "Unmodified")) %>% 
  mutate(
    night_c = as.vector(scale(night2, scale=FALSE)), 
    temp_c  = as.vector(scale(Temp_C, scale=FALSE)), 
    count_stb_c = as.vector(scale(count_stb, scale=FALSE)),
    count_pipe_c = as.vector(scale(count_pipe, scale=FALSE)) # <--- ADDED THIS LINE
  )

# ==============================================================================
# 3. SUPPORTING INFORMATION (SI) FIGURES
# ==============================================================================

# --- Figure S1: Correlation Matrix ---
cor_data <- df %>% 
  filter(species == "Neomysis integer") %>% 
  dplyr::select(Temp_C, night2, Julian_Day, Turbidity_FTU, count_stb, Trap_Type) %>% 
  rename(Temp = "Temp_C", Night = "night2", Turbidity = "Turbidity_FTU", STB = "count_stb", Trap = "Trap_Type") %>%
  na.omit() 

p_s1 <- ggpairs(
  cor_data, 
  columns = 1:5,                 
  mapping = aes(color = Trap, alpha = 0.5), 
  lower = list(continuous = wrap("points", size = 0.5)),
  upper = list(continuous = wrap("cor", size = 2.5)), 
  diag = list(continuous = wrap("densityDiag", alpha = 0.5))
) + 
  theme_custom() +
  theme(panel.grid.major = element_line(colour = "grey90", linewidth = 0.2)) 

ggsave("Graphs/Figure_S1_Correlation.svg", p_s1, width = 170, height = 170, units = "mm")

# --- Figure S2: Benthic Stickleback Size (Histogram) ---
fishtrap2 <- fishtrap %>% 
  distinct(Trap_ID, .keep_all = TRUE) %>% 
  select(Date_Out, Trap_ID, Bay_Name) %>%
  na.omit()

fishtrap2$Trap_ID_bay <- str_extract(fishtrap2$Trap_ID, "^[0-9]+[A-D]")
STSsizeFT <- stb_size %>% mutate(Sample.bay = paste0(Trip, Bay))
fishtrap3 <- fishtrap2 %>% 
  distinct(Trap_ID_bay, .keep_all = TRUE) %>% 
  select(Trap_ID_bay, Date_Out)

stssize_join <- left_join(STSsizeFT, fishtrap3, by = c("Sample.bay" = "Trap_ID_bay"))
stssize_join$Date_Outd <- as.Date(stssize_join$Date_Out)
ordered_dates_sts <- format(sort(unique(stssize_join$Date_Outd)), "%b %d")

stssize_join$Date_Outmd <- format(stssize_join$Date_Outd, "%b %d")
stssize_join$Date_Outmd <- factor(stssize_join$Date_Outmd, levels = ordered_dates_sts)

n_labs_sts <- stssize_join %>% 
  filter(!is.na(Date_Outmd)) %>%
  group_by(Date_Outmd) %>% 
  count() %>% 
  mutate(label = paste0("n=", n), x = 80)

fig_s2 <- ggplot(data = subset(stssize_join, !is.na(Date_Outmd)), 
                 aes(x = Length_mm, y = Date_Outmd)) +
  geom_density_ridges(
    stat = "binline", binwidth = 2, scale = 0.9, draw_baseline = TRUE,
    fill = col_stb, alpha = 1.0, size = 0.2
  ) +
  geom_text(
    data = n_labs_sts, aes(x = x, y = Date_Outmd, label = label),
    inherit.aes = FALSE, size = 2, vjust = -0.5
  ) +
  labs(x = "Length (mm)", y = NULL) +
  coord_cartesian(xlim = c(25, 85)) +
  scale_x_continuous(breaks = seq(30, 80, by = 10)) +
  theme_custom()

ggsave("Graphs/Figure_S2_SticklebackSize.svg", fig_s2, width = 85, height = 100, units = "mm")

# --- Figure S3: Environmental Time Series ---
env_sum <- df %>% 
  group_by(Date_Out, Bay_ID) %>% 
  summarise(
    `Temperature (°C)` = mean(Temp_C, na.rm=T), 
    `Salinity (PSU)` = mean(Salinity_PSU, na.rm=T), 
    `Dissolved Oxygen (mg/L)` = mean(DO_mg_L, na.rm=T), 
    `Turbidity (FTU)` = mean(Turbidity_FTU, na.rm=T), 
    `Chlorophyll a (ppb)` = mean(Chl_a_ppb, na.rm=T), 
    .groups="drop"
  ) %>% 
  pivot_longer(
    cols = c(`Temperature (°C)`, `Salinity (PSU)`, `Dissolved Oxygen (mg/L)`, 
             `Turbidity (FTU)`, `Chlorophyll a (ppb)`),
    names_to = "Variable", values_to = "Value"
  )

env_sum$Variable <- factor(env_sum$Variable, levels = c(
  "Temperature (°C)", "Salinity (PSU)", "Dissolved Oxygen (mg/L)", 
  "Turbidity (FTU)", "Chlorophyll a (ppb)"
))

p_s3 <- ggplot(env_sum, aes(x = as.Date(Date_Out), y = Value, color = as.factor(Bay_ID))) +
  geom_line(linewidth = 0.8) + geom_point(size = 2) +
  facet_wrap(~Variable, scales = "free_y", ncol = 3) +
  scale_color_manual(values = wes_palette("FantasticFox1", n = 5, type = "discrete")) +
  scale_x_date(date_labels = "%b %d", date_breaks = "2 weeks") +
  labs(x = "Date", y = NULL, color = "Area") + 
  theme_custom() +
  theme(
    strip.text = element_text(face = "bold", size = 9), 
    axis.text.x = element_text(angle = 45, hjust = 1),  
    legend.position = "bottom", legend.key = element_rect(fill = NA)
  )

ggsave("Graphs/Figure_S3_EnvVariables.svg", p_s3, width = 170, height = 120, units = "mm")

# --- Figure S4: Precision Analysis (CV vs Effort) ---
neo_data <- lighttrap %>%
  filter(species == "Neomysis integer", grepl("Modified", Trap_Type, ignore.case = TRUE)) %>%
  select(count)

stb_data <- lighttrap %>%
  filter(species %in% c("Gasterosteus aculeatus", "Pungitius pungitius")) %>%
  filter(grepl("Unmodified", Trap_Type, ignore.case = TRUE)) %>%
  group_by(Trap_ID) %>%
  summarise(count = sum(count, na.rm = TRUE), .groups = "drop")

get_params <- function(counts) {
  mu <- mean(counts, na.rm = TRUE)
  v  <- var(counts, na.rm = TRUE)
  if(v <= mu || v == 0) { theta <- 100 } else { theta <- (mu^2) / (v - mu) }
  return(list(mu = mu, theta = theta))
}

p_neo <- get_params(neo_data$count)
p_stb <- get_params(stb_data$count)

simulate_hewitt_cv <- function(mu, theta, max_n = 30, n_sims = 10000) {
  results <- data.frame(n = 2:max_n, cv_95 = NA)
  for (i in 1:nrow(results)) {
    n <- results$n[i]
    sim_matrix <- matrix(rnbinom(n * n_sims, mu = mu, size = theta), nrow = n, ncol = n_sims)
    se_vec <- apply(sim_matrix, 2, sd) / sqrt(n)
    cv_vec <- se_vec / mu 
    results$cv_95[i] <- quantile(cv_vec, 0.95, na.rm = TRUE)
  }
  return(results)
}

set.seed(123)
df_plot <- rbind(
  simulate_hewitt_cv(p_neo$mu, p_neo$theta) %>% mutate(Group = "N. integer (Modified)"),
  simulate_hewitt_cv(p_stb$mu, p_stb$theta) %>% mutate(Group = "Sticklebacks (Unmodified)")
)

optimal_n <- df_plot %>%
  group_by(Group) %>%
  arrange(n) %>%
  mutate(slope = cv_95 - lead(cv_95), initial_slope = slope[1]) %>%
  filter(slope < 0.20 * initial_slope) %>%
  slice(1) %>%
  ungroup() %>%
  mutate(label_y = ifelse(grepl("Stickleback", Group), 
                          max(df_plot$cv_95) * 0.40, 
                          max(df_plot$cv_95) * 0.70))

p_s4 <- ggplot(df_plot, aes(x = n, y = cv_95, color = Group)) +
  geom_line(size = 1) + geom_point(size = 2) +
  geom_vline(data = optimal_n, aes(xintercept = n, color = Group), 
             linetype = "longdash", size = 0.6, show.legend = FALSE) +
  geom_text(data = optimal_n, 
            aes(x = n + 0.6, y = label_y, label = paste0("Optimal n = ", n)), 
            hjust = 0, size = 3, fontface = "bold", show.legend = FALSE) +
  scale_color_manual(values = c("N. integer (Modified)" = "#b30000", 
                                "Sticklebacks (Unmodified)" = "#43a2ca")) +
  scale_x_continuous(breaks = seq(0, 30, 5), limits = c(2, 32)) +
  theme_custom() +  
  labs(y = "CV (95th Percentile)", x = "Number of traps per area") +
  theme(
    legend.position = "bottom", legend.title = element_blank(),
    legend.text = element_text(size = 9), legend.key.size = unit(4, "mm"),
    axis.title = element_text(size = 10), axis.text = element_text(size = 9),
    panel.grid.major = element_line(color = "grey92", size = 0.3), 
    panel.grid.minor = element_blank()
  ) +
  guides(color = guide_legend(ncol = 1))

ggsave("Graphs/Figure_S4.svg", p_s4, width = 85, height = 85, units = "mm")

# ==============================================================================
# 4. SUPPORTING INFORMATION (SI) TABLES
# ==============================================================================

m_neo_night <- glmmTMB(count ~ Trap_Type + night_c + (1|Bay_ID/Station), family=nbinom2, data=neo)
m_neo_temp  <- glmmTMB(count ~ Trap_Type + temp_c + (1|Bay_ID/Station), family=nbinom2, data=neo) 
m_pra_night <- glmmTMB(count ~ Trap_Type + night_c + (1|Bay_ID/Station), family=nbinom2, data=pra)
m_pra_temp  <- glmmTMB(count ~ Trap_Type + temp_c + (1|Bay_ID/Station), family=nbinom2, data=pra) 
m_stb_night <- glmmTMB(count ~ Trap_Type + night_c + (1|Bay_ID/Station), family=nbinom2, data=stb_pooled)
m_stb_temp  <- glmmTMB(count ~ Trap_Type + temp_c + (1|Bay_ID/Station), family=nbinom2, data=stb_pooled) 

tab_s1_data <- data.frame(
  Species = c("N. integer", "P. flexuosus", "G. aculeatus + P. pungitius"),
  Trap_type = "Light trap", 
  AIC_Night = c(AIC(m_neo_night), AIC(m_pra_night), AIC(m_stb_night)),
  AIC_Temp = c(AIC(m_neo_temp), AIC(m_pra_temp), AIC(m_stb_temp))
) %>% 
  mutate(
    Delta_AIC = AIC_Temp - AIC_Night,
    Selected_Driver = ifelse(Delta_AIC > 2, "Night Duration", "Inconclusive") 
  )

border_top_double <- fp_border(color = "black", width = 4, style = "double") 
border_inner_thin <- fp_border(color = "black", width = 1, style = "solid")

ft_s1 <- flextable(tab_s1_data) %>%
  set_header_labels(
    Species = "Species/Group", Trap_type = "Trap type",
    AIC_Night = "Model: Night duration (AIC)", AIC_Temp = "Model: Temperature (AIC)",
    Delta_AIC = "ΔAIC", Selected_Driver = "Selected Driver"
  ) %>%
  colformat_double(j = c("AIC_Night", "AIC_Temp", "Delta_AIC"), digits = 1) %>%
  bold(part = "header") %>% align(align = "left", part = "all") %>%
  border_remove() %>% hline_top(part = "header", border = border_top_double) %>%
  hline_bottom(part = "header", border = border_inner_thin) %>%
  hline_bottom(part = "body", border = border_inner_thin) %>% autofit()
print(ft_s1)

# --- Table S2: Combined Mysid Model ---
mys_comb <- df %>% filter(species %in% c("Neomysis integer", "Praunus flexuosus"), Trap_Type %in% c("Modified net", "Unmodified")) %>% 
  mutate(night_c=scale(night2, scale=F))
m_s2 <- glmmTMB(count ~ species * Trap_Type + species * night_c + (1|Bay_ID/Station), family=nbinom2, data=mys_comb)
tab_model(m_s2, show.se=T, show.stat=T, title="Table S2")

# --- Table S3: Stickleback Species Interaction ---
stb_indiv <- df %>% filter(species %in% c("Gasterosteus aculeatus", "Pungitius pungitius"), Trap_Type %in% c("Modified net", "Unmodified")) %>% 
  mutate(night_c=scale(night2, scale=F))
m_s3 <- glmmTMB(count ~ species * night_c + species * temp_c+ (1|Bay_ID/Station), family=nbinom2, data=stb_indiv)
tab_model(m_s3, show.se=T, show.stat=T, title="Table S3")

# --- Table S4: Prevalence Summary ---
lt_stats <- df %>%
  filter(Trap_Type == "Modified net", species %in% c("Neomysis integer", "Praunus flexuosus")) %>%
  filter(!is.na(count)) %>% group_by(species) %>%
  summarise(
    Trap_type = "Light trap (Modified)", Total_traps = n(),
    Prevalence = sum(count > 0), Freq_occurrence = (sum(count > 0) / n()) * 100,
    Individuals = sum(count), .groups = "drop"
  ) %>%
  mutate(Percent_catch = (Individuals / sum(Individuals)) * 100)

bt_stats <- fishtrap %>%
  filter(TRIP != 1, species %in% c("Gasterosteus aculeatus", "Pungitius pungitius")) %>%
  filter(!is.na(count)) %>% group_by(species) %>%
  summarise(
    Trap_type = "Benthic trap", Total_traps = n(), 
    Prevalence = sum(count > 0), Freq_occurrence = (sum(count > 0) / n()) * 100,
    Individuals = sum(count), .groups = "drop"
  ) %>%
  mutate(Percent_catch = (Individuals / sum(Individuals)) * 100)

tab_s4_data <- bind_rows(lt_stats, bt_stats) %>%
  select(Trap_type, species, Total_traps, Prevalence, Freq_occurrence, Individuals, Percent_catch)

ft_s4 <- flextable(tab_s4_data) %>%
  set_header_labels(
    Trap_type = "Trap type", species = "Species", Total_traps = "Total traps (n)",
    Prevalence = "Prevalence (n traps)", Freq_occurrence = "Freq. occurrence (%)",
    Individuals = "Individuals (n)", Percent_catch = "Percent catch (%)"
  ) %>%
  colformat_double(j = c("Freq_occurrence", "Percent_catch"), digits = 1) %>%
  colformat_double(j = c("Total_traps", "Prevalence", "Individuals"), digits = 0) %>%
  bold(part = "header") %>% align(align = "left", part = "all") %>% border_remove() %>%
  hline_top(part = "header", border = border_top_double) %>%
  hline_bottom(part = "header", border = border_inner_thin) %>%
  hline_bottom(part = "body", border = border_inner_thin) %>% autofit()
print(ft_s4)

# --- Table S5: Spatial Autocorrelation ---
calc_spatial_autocorr <- function(model, data, species_label) {
  set.seed(42) 
  res <- simulateResiduals(model, plot = FALSE)
  res_agg <- recalculateResiduals(res, group = data$Station_comb)
  
  spatial_meta <- data %>%
    group_by(Station_comb) %>%
    summarise(Bay_ID = first(Bay_ID), Long_DD = first(Long_DD), Lat_DD = first(Lat_DD), .groups = "drop") %>%
    arrange(Station_comb) 
  
  spatial_meta$residuals <- res_agg$scaledResiduals
  results_list <- list()
  bays <- sort(unique(spatial_meta$Bay_ID))
  
  for(b in bays) {
    bay_data <- spatial_meta %>% filter(Bay_ID == b)
    if(nrow(bay_data) < 3) next 
    test <- testSpatialAutocorrelation(simulationOutput = bay_data$residuals, x = bay_data$Long_DD, y = bay_data$Lat_DD, plot = FALSE)
    results_list[[b]] <- data.frame(Species = species_label, Area = b, Morans_I = test$statistic[1], p_value = test$p.value)
  }
  return(bind_rows(results_list))
}

df_neo_spatial <- calc_spatial_autocorr(m_neo_night, neo, "N. integer")
df_pra_spatial <- calc_spatial_autocorr(m_pra_night, pra, "P. flexuosus")
tab_s5_data <- bind_rows(df_neo_spatial, df_pra_spatial)

ft_s5 <- flextable(tab_s5_data) %>%
  set_header_labels(Species = "Species", Area = "Area", Morans_I = "Moran's I", p_value = "p-value") %>%
  colformat_double(j = "Morans_I", digits = 2) %>% colformat_double(j = "p_value", digits = 2) %>%
  bold(part = "header") %>% align(align = "left", part = "all") %>% border_remove() %>%
  hline_top(part = "header", border = fp_border(width = 4, style = "double")) %>%
  hline_bottom(part = "header", border = fp_border(width = 1)) %>%
  hline_bottom(part = "body", border = fp_border(width = 1)) %>% autofit()
print(ft_s5)

# --- Table S6: Catch Composition ---
species_order <- lighttrap %>% group_by(species) %>% summarise(Total = sum(count, na.rm = TRUE)) %>% arrange(desc(Total)) %>% pull(species)

tab_s6_data <- lighttrap %>%
  group_by(Date_Out, Bay_ID, Trap_Type, species) %>%
  summarise(Count = sum(count, na.rm = TRUE), .groups = "drop") %>%
  pivot_wider(names_from = species, values_from = Count, values_fill = 0) %>%
  arrange(as.Date(Date_Out), Bay_ID) %>%
  select(Date_Out, Bay_ID, Trap_Type, all_of(species_order))

ft_s6 <- flextable(tab_s6_data) %>%
  set_header_labels(Date_Out = "Date", Bay_ID = "Area", Trap_Type = "Trap type") %>%
  italic(part = "header", j = ~ . -Date_Out -Bay_ID -Trap_Type) %>% 
  bold(part = "header") %>% align(align = "left", part = "all") %>% border_remove() %>%
  hline_top(part = "header", border = border_top_double) %>%
  hline_bottom(part = "header", border = border_inner_thin) %>%
  hline_bottom(part = "body", border = border_inner_thin) %>% autofit()
print(ft_s6)

# --- Table S7: Detailed Mysid Selection ---
tab_model(
  m_neo_night, m_neo_temp, m_pra_night, m_pra_temp,
  transform = NULL, show.se = TRUE, show.ci = FALSE,
  dv.labels = c("N. integer (Night)", "N. integer (Temp)", "P. flexuosus (Night)", "P. flexuosus (Temp)"),
  string.est = "Log Mean", string.se = "SE", digits = 2, show.stat = TRUE, show.p = TRUE, title = "Table S7"
)

# --- Table S8: Benthic Model Selection ---
m_ft_full   <- glmmTMB(count ~ temp_c + night_c + (1|Bay_ID/Station), family=nbinom2, data=ft_sts)
m_ft_night  <- glmmTMB(count ~ night_c + (1|Bay_ID/Station), family=nbinom2, data=ft_sts)
m_ft_temp   <- glmmTMB(count ~ temp_c + (1|Bay_ID/Station), family=nbinom2, data=ft_sts)

tab_model(
  m_ft_full, m_ft_night, m_ft_temp,
  dv.labels = c("Full Model", "Night only", "Temperature only"),
  show.ci = FALSE, show.se = TRUE, show.stat = TRUE, collapse.se = FALSE,      
  string.est = "IRR", string.se = "SE", show.aic = TRUE, title="Table S8"
)

# ==============================================================================
# 5. MAIN TEXT TABLES & MODELS
# ==============================================================================

# --- Table 1: Stickleback Model Selection ---
m_stb_full <- glmmTMB(count ~ Trap_Type + night_c + temp_c + (1|Bay_ID/Station), family=nbinom2, data=stb_pooled)
tab_model(m_stb_full, m_stb_night, m_stb_temp, dv.labels=c("Full Model", "Night Only", "Temp Only"), show.aic=T, show.se=T, show.stat=T, p.style="numeric", title="Table 1")

# --- Table 2: Mysid Models & Biotic Interaction ---
m_neo_stb <- glmmTMB(count ~ count_stb_c + night_c + (1|Bay_ID/Station), family=nbinom2, data=neo)
tab_model(m_neo_night, m_pra_night, m_neo_stb, dv.labels=c("Neomysis (Mod1)", "Praunus (Mod2)", "Neomysis (Mod1b)"), show.ci = F, show.se=T, show.stat=T, p.style="numeric", title="Table 2")

# Pipefish control model:
m_neo_pipe <- glmmTMB(count ~ count_pipe_c + night_c + (1|Bay_ID/Station), family=nbinom2, data=neo)
summary(m_neo_pipe)
# Updated Table output to include the pipefish model:
tab_model(
  m_neo_night, m_neo_stb, m_neo_pipe, 
  dv.labels=c("Neomysis (Base)", "Neomysis (Stickleback)", "Neomysis (Pipefish Control)"), 
  show.ci = F, show.se=T, show.stat=T, p.style="numeric", title="Table 2"
)

# ==============================================================================
# 6. MAIN TEXT FIGURES
# ==============================================================================

# --- FIGURE 4: N. integer ---
mean_night_neo <- mean(neo$night2, na.rm = TRUE)
mean_stb_neo   <- mean(neo$count_stb, na.rm = TRUE)
mean_night_pra <- mean(pra$night2, na.rm = TRUE)

theme_clean <- theme_custom() + theme(plot.title = element_blank(), axis.title = element_text(size = 9, face = "bold"), axis.text = element_text(size = 8))

p4a <- plot_model(m_neo_night, type = "pred", terms = "Trap_Type", colors = col_neo) + labs(tag = "A", y = expression(paste("Predicted count (", italic("N. integer"), ")")), x = "Trap type", title = "") + theme_clean
p4b_raw <- plot_model(m_neo_night, type = "pred", terms = "night_c [all]", colors = col_neo)
p4b_raw$data$x <- p4b_raw$data$x + mean_night_neo  
p4b <- p4b_raw + labs(tag = "B", y = "", x = "Night duration (h)", title = "") + theme_clean
p4c_raw <- plot_model(m_neo_stb, type = "pred", terms = "count_stb_c [all]", colors = col_neo)
p4c_raw$data$x <- p4c_raw$data$x + mean_stb_neo    
p4c <- p4c_raw + labs(tag = "C", y = "", x = "Stickleback abundance (n)", title = "") + theme_clean

p4_final <- p4a + p4b + p4c + plot_layout(nrow = 2)
ggsave("Graphs/Figure_4_Neomysis_RawScale.svg", p4_final, width = 170, height = 160, units = "mm")

# --- FIGURE 5: P. flexuosus ---
p5a <- plot_model(m_pra_night, type = "pred", terms = "Trap_Type", colors = col_pra) + labs(tag = "A", y = expression(paste("Predicted count (", italic("P. flexuosus"), ")")), x = "Trap type", title = "") + theme_clean
p5b_raw <- plot_model(m_pra_night, type = "pred", terms = "night_c [all]", colors = col_pra)
p5b_raw$data$x <- p5b_raw$data$x + mean_night_pra  
p5b <- p5b_raw + labs(tag = "B", y = "", x = "Night duration (h)", title = "") + theme_clean

p5_final <- p5a + p5b + plot_layout(nrow = 1)
ggsave("Graphs/Figure_5_Praunus_RawScale.svg", p5_final, width = 170, height = 80, units = "mm")

# --- FIGURE 6: BENTHIC TRAP G. aculeatus ---
mean_temp_ft  <- mean(ft_sts$Temp_C, na.rm = TRUE)
mean_night_ft <- mean(ft_sts$night2, na.rm = TRUE)

p6a_raw <- plot_model(m_ft_temp, type = "pred", terms = "temp_c [all]", colors = col_stb)
p6a_raw$data$x <- p6a_raw$data$x + mean_temp_ft  
p6a <- p6a_raw + labs(tag = "A", y = expression(paste("Predicted count (", italic("G. aculeatus"), ")")), x = "Temperature (°C)", title = "") + theme_clean
p6b_raw <- plot_model(m_ft_night, type = "pred", terms = "night_c [all]", colors = col_stb)
p6b_raw$data$x <- p6b_raw$data$x + mean_night_ft 
p6b <- p6b_raw + labs(tag = "B", y = "", x = "Night duration (h)", title = "") + theme_clean

fig6_final <- p6a + p6b + plot_layout(nrow = 1)
ggsave("Graphs/Figure_6_Benthic_RawScale.svg", fig6_final, width = 170, height = 80, units = "mm")

# --- FIGURE 7: MYSID SIZE STRUCTURE ---
plot_data <- mysid_size %>%
  left_join(lighttrap %>% distinct(Trap_ID, .keep_all = TRUE), by = c("Sample" = "Trap_ID")) %>%
  filter(!is.na(Date_Out), Trap_Type == "Modified net") %>%
  mutate(
    Date_Outd = as.Date(Date_Out),
    Date_Outmd = factor(format(Date_Outd, "%b %d"), levels = format(sort(unique(Date_Outd)), "%b %d")),
    Species = case_when(grepl("integer", Species) ~ "Neomysis integer", TRUE ~ "Praunus flexuosus")
  )

n_labels <- plot_data %>% count(Date_Outmd, Species) %>% complete(Date_Outmd, Species, fill = list(n = 0)) %>%
  mutate(label = paste0("n=", n), x_pos = ifelse(Species == "Neomysis integer", 0, 26), text_col = ifelse(Species == "Neomysis integer", col_neo, col_pra))

fig7 <- ggplot(plot_data, aes(x = Length_mm, y = Date_Outmd, fill = Species)) +
  geom_density_ridges(aes(height = ..count..), stat = "binline", bins = 45, scale = 0.95, alpha = 0.7, draw_baseline = TRUE, color = "black", size = 0.2) +
  geom_text(data = n_labels, aes(x = x_pos, y = Date_Outmd, label = label), color = n_labels$text_col, size = 3, fontface = "bold", vjust = -1.5, inherit.aes = FALSE) +
  scale_fill_manual(values = c("Neomysis integer" = col_neo, "Praunus flexuosus" = col_pra), labels = c(expression(italic("N. integer")), expression(italic("P. flexuosus")))) +
  scale_y_discrete(expand = expansion(mult = c(0.01, 0.15))) + coord_cartesian(xlim = c(-2, 30)) +
  theme_custom() + labs(x = "Length (mm)", y = "") +
  theme(legend.position = "bottom", legend.key.size = unit(3, "mm"), legend.text = element_text(size = 9), legend.margin = margin(t = -5), axis.text.y = element_text(vjust = 0), plot.margin = margin(5, 5, 5, 5, "mm")) + guides(fill = guide_legend(title = NULL))

ggsave("Graphs/Figure_7_MysidSize_Final.svg", fig7, width = 85, height = 140, units = "mm")

# ==============================================================================
# 7. POWER ANALYSIS (Figures 8 & 9)
# ==============================================================================

simulate_scenario <- function(n_traps, theta, sd_bay, intercept, effect_size=0.5, n_bays=4, n_sims=500) {
  slope <- log(effect_size) 
  significant_runs <- 0; valid_runs <- 0
  for(i in 1:n_sims) {
    bays  <- rep(paste0("Bay", 1:n_bays), each = n_traps * 2)
    years <- rep(rep(c("Y1", "Y2"), each = n_traps), times = n_bays)
    bay_offsets <- rnorm(n_bays, mean = 0, sd = sd_bay)
    names(bay_offsets) <- paste0("Bay", 1:n_bays)
    mus <- exp(intercept + ifelse(years == "Y2", slope, 0) + bay_offsets[bays])
    counts <- rnbinom(n = length(years), mu = mus, size = theta)
    df_sim <- data.frame(Bay = bays, Year = factor(years, levels = c("Y1", "Y2")), count = counts)
    m_sim <- try(suppressWarnings(glmmTMB(count ~ Year + (1|Bay), family = nbinom2, data = df_sim)), silent = TRUE)
    if(!inherits(m_sim, "try-error") && m_sim$fit$convergence == 0) {
      valid_runs <- valid_runs + 1
      p_val <- summary(m_sim)$coefficients$cond["YearY2", "Pr(>|z|)"]
      if(!is.na(p_val) && p_val < 0.05) significant_runs <- significant_runs + 1
    }
  }
  if(valid_runs == 0) return(0)
  return(significant_runs / valid_runs)
}

simulate_distribution <- function(n_traps, theta, sd_bay, intercept, effect_size=0.5, n_bays=4, n_sims=1000) {
  slope <- log(effect_size)
  results <- data.frame(Trial = integer(), Group = character(), Mean_Count = numeric())
  for(i in 1:n_sims) {
    bays  <- rep(paste0("Bay", 1:n_bays), each = n_traps * 2)
    years <- rep(rep(c("Y1", "Y2"), each = n_traps), times = n_bays)
    bay_offsets <- rnorm(n_bays, mean = 0, sd = sd_bay)
    mus <- exp(intercept + ifelse(years == "Y2", slope, 0) + bay_offsets[rep(1:n_bays, each=n_traps*2)])
    counts <- rnbinom(n = length(years), mu = mus, size = theta)
    mean_y1 <- mean(counts[years == "Y1"]); mean_y2 <- mean(counts[years == "Y2"])
    results <- rbind(results, data.frame(Trial = i, Group = "Reference", Mean_Count = mean_y1), data.frame(Trial = i, Group = "Impact (-50%)", Mean_Count = mean_y2))
  }
  return(results)
}

mysid_power_data <- df %>% filter(species == "Neomysis integer", Trap_Type == "Modified net") %>% droplevels()
m_ref_neo <- glmmTMB(count ~ night_c + (1|Bay_ID), data = mysid_power_data, family = nbinom2)
theta_neo  <- sigma(m_ref_neo)
sd_bay_neo <- attr(VarCorr(m_ref_neo)$cond$Bay_ID, "stddev")
if(is.null(sd_bay_neo) || is.na(sd_bay_neo)) sd_bay_neo <- 0.5 
standard_int <- log(5) 

stb_power_data <- stb_pooled %>% filter(Trap_Type == "Unmodified") %>% droplevels()
m_ref_stb <- glmmTMB(count ~ night_c + (1|Bay_ID), data = stb_power_data, family = nbinom2)
theta_stb  <- sigma(m_ref_stb)
sd_bay_stb <- attr(VarCorr(m_ref_stb)$cond$Bay_ID, "stddev")
if(is.null(sd_bay_stb) || is.na(sd_bay_stb)) sd_bay_stb <- 0.5

traps_vec <- c(5, 10, 15, 20); declines  <- c(0.5, 0.4, 0.3, 0.2, 0.1); increases <- c(1.5, 2.0, 3.0, 5.0)      

res_neo_A <- sapply(traps_vec, function(n) simulate_scenario(n, theta_neo, sd_bay_neo, standard_int, 0.5))
res_neo_Sens <- sapply(declines, function(d) simulate_scenario(10, theta_neo, sd_bay_neo, standard_int, d))
res_neo_D <- sapply(increases, function(i) simulate_scenario(10, theta_neo, sd_bay_neo, standard_int, i))
dist_neo <- simulate_distribution(n_traps=10, theta_neo, sd_bay_neo, standard_int, 0.5)

res_stb_A <- sapply(traps_vec, function(n) simulate_scenario(n, theta_stb, sd_bay_stb, standard_int, 0.5))
res_stb_Sens <- sapply(declines, function(d) simulate_scenario(10, theta_stb, sd_bay_stb, standard_int, d))
res_stb_D <- sapply(increases, function(i) simulate_scenario(10, theta_stb, sd_bay_stb, standard_int, i))
dist_stb <- simulate_distribution(n_traps=10, theta_stb, sd_bay_stb, standard_int, 0.5)

make_line_plot <- function(x_data, y_data, title, xlabel, color_val, type="line") {
  df <- data.frame(x = x_data, y = y_data)
  p <- ggplot(df, aes(x, y)) + geom_hline(yintercept = 0.8, linetype = "dashed", color = "grey50") + geom_line(color = color_val, size = 1) + geom_point(color = color_val, size = 2) + labs(title = title, y = "Power", x = xlabel) + theme_custom() + scale_y_continuous(limits = c(0, 1), breaks = seq(0, 1, 0.2))
  if(type == "percent") { p <- p + scale_x_reverse() }
  return(p)
}

make_density_plot <- function(dist_data, color_val, title) {
  ggplot(dist_data, aes(x = Mean_Count, fill = Group)) + geom_density(alpha = 0.6, color = NA) + scale_fill_manual(values = c("Reference" = "grey70", "Impact (-50%)" = color_val)) + labs(title = title, x = "Simulated mean catch (n=10)", y = "Density") + theme_custom() + theme(legend.position = c(0.7, 0.8), legend.background = element_rect(fill = "white", color = NA), legend.key.size = unit(3, "mm"))
}

p8a <- make_line_plot(traps_vec, res_neo_A, "", "Sample size (n)", col_neo) + labs(tag="A")
p8b <- make_density_plot(dist_neo, col_neo, "") + labs(tag="B")
p8c <- make_line_plot((1-declines)*100, res_neo_Sens, "", "Decline magnitude (%)", col_neo, type="percent") + labs(tag="C")
p8d <- make_line_plot(increases, res_neo_D, "", "Fold increase (x)", col_neo) + labs(tag="D", y="") + scale_x_continuous(breaks=increases, labels=paste0(increases,"x"))

fig8 <- (p8a + p8b) / (p8c + p8d)
ggsave("Graphs/Figure_8_MysidPower.svg", fig8, width=170, height=150, units="mm")

p9a <- make_line_plot(traps_vec, res_stb_A, "", "Sample Size (n)", col_stb) + labs(tag="A")
p9b <- make_density_plot(dist_stb, col_stb, "") + labs(tag="B")
p9c <- make_line_plot((1-declines)*100, res_stb_Sens, "", "Decline Magnitude (%)", col_stb, type="percent") + labs(tag="C")
p9d <- make_line_plot(increases, res_stb_D, "", "Fold Increase (x)", col_stb) + labs(tag="D", y="") + scale_x_continuous(breaks=increases, labels=paste0(increases,"x"))

fig9 <- (p9a + p9b) / (p9c + p9d)
ggsave("Graphs/Figure_9_SticklebackPower.svg", fig9, width=170, height=150, units="mm")

print("--- POWER ANALYSIS COMPLETE ---")
```
